# GALA: A Unified Landmark-Free Framework for Coarse-to-Fine Spatial Alignment Across Resolutions and Modalities in Spatial Transcriptomics

**DOI:** 10.64898/2025.11.29.691288

**Authors:** Tao Ding, Pengcheng Zeng

## Abstract

Spatial transcriptomics alignment is challenged by technical variations, including geometric distortions from tissue preparation and platform-driven differences in resolution and modality. These issues create diverse alignment scenarios, from matched and mismatched resolutions to cross-modality integration, while partial tissue coverage further complicates the task. To overcome these limitations, we introduce GALA (Genetic Algorithm–guided Large Deformation Alignment), a unified, landmark-free framework that couples global affine transformation and local diffeomorphic deformation within a single optimisation. Its modality-aware rasterisation harmonises transcriptomic and histological data into a shared grid, enabling landmark-free, multimodal alignment across resolutions and modalities. Evaluated on diverse human and mouse datasets, GALA outperforms existing methods in accuracy, computational efficiency, and biological interpretability for both complete and partial tissue alignment.

## 1. Introduction

Spatial transcriptomics (ST) technologies enable high-throughput mapping of gene expression within intact tissue sections, providing detailed insights into cellular organisation, tissue architecture and disease-associated microenvironments [1, 2, 3]. Contemporary platforms, including 10x Visium [4], Xenium [5] and MERFISH [6], span a wide range of spatial resolutions, from multi-cellular spots (spot-level) to single-cell or subcellular precision (cell-level), and frequently integrate transcriptomic measurements with histological images. While this diversity expands analytical possibilities, it also introduces substantial variability across tissues, samples and platforms, making direct spatial comparison and integration challenging [7, 8, 9]. Geometric distortions arising during tissue preparation, such as rotation, shifting, stretching and local warping, further complicate alignment, even between adjacent sections from the same specimen [10, 11, 12]. Correcting these transformations is essential to preserve tissue morphology and establish accurate spatial correspondence, enabling downstream applications such as cross-sample integration, three-dimensional reconstruction, deconvolution and spatial domain annotation [8, 13, 14, 15]. For example, modelling cell–cell regulatory interactions or perturbation effects [16, 17, 18] relies on coherent spatial coordinates to interpret functional relationships. Geometric alignment therefore constitutes a foundational step for spatially informed biological inference.

Beyond geometric distortion, ST datasets often differ in spatial resolution, gene coverage and molecular sensitivity across platforms [3, 19]. These discrepancies complicate the integration of measurements across resolutions, from single-cell transcriptomes to multi-cellular spot profiles, as well as across modalities, including histology and transcriptomics. In practice, ST alignment encompasses three common scenarios: matched-resolution (spot-to-spot or cell-to-cell), mismatched-resolution (cell-to-spot), and cross-modality (transcriptomics-to-histology). Each scenario imposes unique methodological constraints, influenced by factors such as spatial resolution, tissue integrity, and the availability of complementary histological context.

Spot-to-spot alignment is typically performed using rigid or affine transformations, and in some cases smooth non-rigid deformations informed by spatial features and gene expression [7, 8], suitable when each spot aggregates multiple cells and local variation is smoothed. Representative methods include PASTE [13], STAligner [20] and GPSA [21], which perform well under matched resolution and full-tissue overlap but struggle with partial coverage, cross-platform differences, or large-scale single-cell data. Cell-to-cell alignment requires non-rigid methods to capture fine anatomical variability [3, 12]; methods such as STalign [12] estimate diffeomorphic deformation fields but need landmark-based pre-alignment and are sensitive to large displacements or missing tissue. Coarse-to-fine frameworks like ST-GEARS [22] combine global affine transformation and local refinement, improving performance in complex misalignments but mainly for matched-resolution data. Mismatched-resolution alignment, e.g., mapping single-cell profiles to spot-level data, enables cell-type annotation and spatial reconstruction [23, 24], though methods like SLAT [25] rely on sparse correspondences and have limited scalability. Tissue morphology, e.g., H&E images, can aid alignment [26]; GPSA [21] and PASTE2 [27] incorporate image features but remain spot-level. Cross-modality alignment, such as transcriptomics-to-histology [12, 28], is challenging due to unshared features and differing distributions, with existing approaches often requiring manual landmarks (e.g., STalign). Incomplete tissue coverage further complicates alignment [29], as most methods assume full overlap, with only limited probabilistic handling in PASTE2 [27].

Despite rapid methodological advances in alignments, existing approaches remain fragmented across problem settings. High-resolution frameworks such as CAST [30] often suffer from substantial memory consumption due to dense point-wise matching. Other approaches, including SPACEL [31] and SPIRAL [32], depend heavily on pre-existing cell annotations or cluster-aware information, limiting applicability in fully unsupervised scenarios. Moreover, many methods assume substantial tissue overlap or matched resolutions, reducing robustness in cross-platform or partially overlapping datasets [7, 8, 33]. Collectively, these limitations highlight the need for a unified, scalable alignment approach capable of handling diverse platforms, spatial resolutions, and modalities, while remaining robust to missing or partially overlapping tissue.

To address these challenges, we introduce *GALA* (**G**enetic **A**lgorithm–guided **L**arge Deformation **A**lignment), a general and fully automated framework for multimodal spatial alignment. GALA combines a modality-aware rasterisation module that projects heterogeneous transcriptomic and histological inputs onto a shared spatial grid. It then employs a coupled optimisation strategy to jointly estimates global affine transformation and local diffeomorphic deformation under a single objective function. This design enables accurate alignment across matched- and mismatched-resolution settings (spot-to-spot, cell-to-cell and cell-to-spot) as well as transcriptomics-to-histology alignment, while naturally supporting partial tissue overlap without manual landmarks. Across human and mouse datasets from Visium, MERFISH and Xenium, GALA delivers accurate, efficient and biologically coherent alignments, facilitating downstream comparative analyses and multi-sample atlas construction.

## 2. Materials and Methods

### 2.1. Overview of the GALA framework

GALA is a unified, landmark-free framework for spatial transcriptomics alignment across resolutions and modalities. It jointly corrects large-scale geometric distortions and fine-scale local variations, while flexibly integrating transcriptomic and histological information when available. As illustrated in Fig. 1a, the workflow comprises four components: Input, Multimodal rasterisation, Coupled alignment process, and Output. The input contains source (to be aligned) and target datasets, each potentially multimodal comprising spatial coordinates, gene expression, and histological images (e.g., H&E staining, assumed to be available). To enable efficient and unified processing, heterogeneous transcriptomic measurements from informative genes and histological images are converted into structured, image-like grids, providing a common representation for efficient joint processing. The coupled alignment process consists of three elements: a global affine operator *A* that captures large-scale linear transformations such as rotation, translation, scaling, and reflection, while identifying the overlapping region between source and target sections; a local diffeomorphic mapping *ϕ*^*v*^ that models smooth nonlinear deformations; and a probabilistic matching term *P*_*M*_ that distinguishes reliably matched areas within the overlapping region. The output stage produces the aligned source data, and for illustration, includes alignment results for a representative gene. Fig. 1b summaries the overall objective structure and optimisation flow, while Fig. 1c highlights the range of alignment scenarios supported by GALA, including matched- and mismatched-resolution alignment, cross-modality alignment, and partial-overlap cases.

**Fig. 1.**
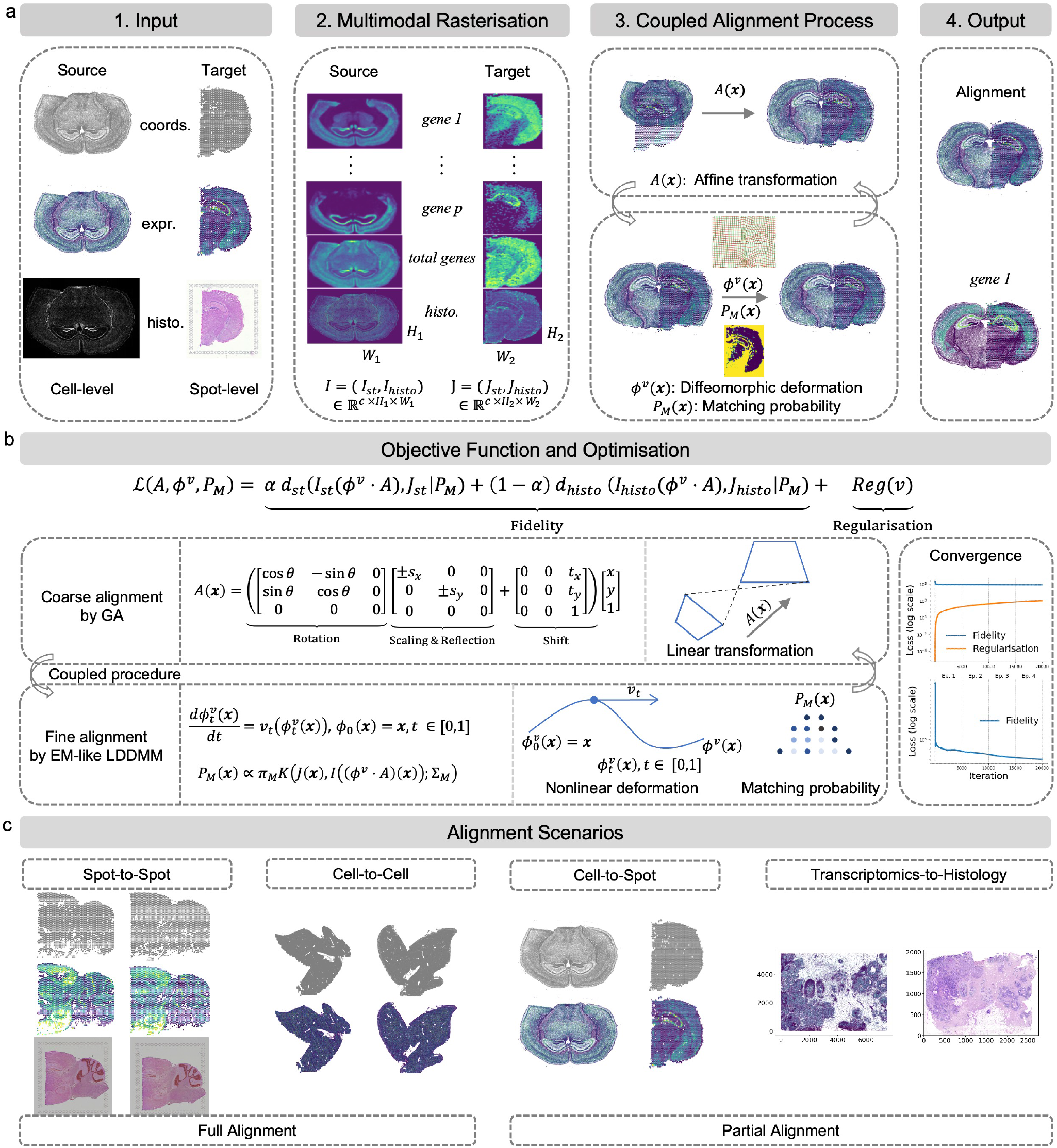
Overview of the GALA framework and its application scenarios. **a** Workflow of GALA. The multimodal source (to be aligned) and target datasets, comprising spatial coordinates (‘coords.’), gene expression (‘expr.’), and histological images (‘histo.’), potentially at different resolutions, are rasterised into multi-channel tensors, *I* = (*I*_st_, *I*_histo_) and *J* = (*J*_st_, *J*_histo_), by concatenating the transcriptomic and histology channels. The coupled alignment jointly estimates a global affine transformation *A*, a local diffeomorphic mapping *ϕ*^*v*^, and a probabilistic matching field *P*_*M*_, yielding the final aligned output. An example gene alignment is shown. **b** Schematic of the unified objective function and optimisation. The objective combines transcriptomic and histological fidelity terms (weighted by *α* and 1 − *α*) with a regularisation on *v*. Fidelity is measured by a weighted squared error *d*(*I, J* | *P*_*M*_) = ∑***x***_∈Ω_ *P*_*M*_ ∥*I* − *J*∥^2^, where the matching probability *P*_*M*_ modulates the contribution of each spatial location. The global stage estimates *A* via a GA, while the local stage jointly refines *ϕ*^*v*^ and *P*_*M*_ through an EM-like LDDMM update, guided by a Gaussian posterior with prior *π*_*M*_ and covariance ∑_*M*_ . Convergence of fidelity and regularisation across global–local iterations is shown. (The bottom one only focuses the fidelity to better visualise its convergence trend). **c** GALA supports a wide range of alignment tasks, including matched-resolution alignment (spot-to-spot and cell-to-cell), mismatched-resolution alignment (cell-to-spot or spot-to-cell), cross-modality alignment (e.g., transcriptomics-to-histology), and partial alignment with incomplete tissue coverage.

### 2.2. Data preprocessing and rasterisation

All datasets used in this study are preprocessed according to quantitative quality-control criteria, including standard procedures such as cell filtering, gene selection, and log-transformation. Both the raw and preprocessed datasets, together with access links, are described in Section 5 and are also provided in the accompanying `GALA` code package.

To integrate transcriptomic and histological information within a unified optimisation framework, GALA represents all data modalities as co-registered raster tensors. Each dataset is projected onto a regular two-dimensional grid Ω with spacing *dx* and dimensions *H × W*, onto which transcriptomic or histological measurements are rasterised. Let ***x***_*i*_ ∈ ℝ^2^ denote the coordinates of spatial anchors (i.e., cells or spots) for *i* = 1, …, *n* and ***y***_*i*_ ∈ ℝ^*p*^ their corresponding gene expression vectors. The rasterised transcriptomic tensor *I*_st_(***x***) is defined as

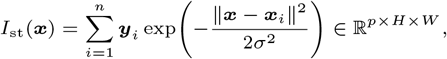

which smooths expression vectors over spatial coordinates using Gaussian kernels, thereby encoding both molecular intensity and spatial proximity. The Gaussian width *σ* is set to 2*dx* by default and can be adjusted to match the spatial characteristics of each dataset: larger values suppress noise through stronger smoothing, whereas smaller values preserve fine-grained spatial patterns (see examples in Supplementary Section S1.1). A sensitivity analysis of *dx* is provided in Supplementary Section S2.1.3. When datasets differ in spatial scale (e.g., cell-level versus spot-level), the Gaussian kernel widths can be tuned independently. This ensures that datasets with mismatched resolutions are rasterised onto grids with comparable effective resolutions, enabling cross-resolution alignment (e.g., cell-to-spot). Moreover, the rasterised representation substantially reduces computational cost by operating on downsampled raster signals rather than high-dimensional molecular matrices. In this study, we typically set *dx* ≈ 1 unit for spot-level data (corresponding to low-resolution scale) and *dx* = 30 *µ*m for cell-level data, balancing spatial fidelity and computational efficiency. To stabilise regions with weak or noisy expression, an additional channel recording the total expression across all genes is included, yielding the final transcriptomic tensor *I*_st_ ∈ ℝ^(*p*+1)*×H×W*^ .

When histological images are available, they are rasterised in parallel onto the same grid used for transcriptomics. Because each image pixel is spatially registered to the transcriptomics, the resulting histological raster is naturally co-aligned with the transcriptomic tensor, enabling a unified multimodal representation for subsequent alignment. Specifically, raw RGB images are first converted to greyscale, and structural contrast is enhanced using gradient magnitude filters (e.g., the Sobel operator [34]). Gaussian smoothing is then applied to generate a single-channel histological raster tensor, *I*_histo_ ∈ ℝ^1*×H×W*^ . Alternatively, RGB images can be processed channel-wise to obtain *I*_histo_ ∈ ℝ^3*×H×W*^ ; however, gradient-based greyscale maps are used by default due to their robust structural representation and reduced sensitivity to staining variation.

Thus, each dataset is ultimately represented by a raster tensor *I* ∈ ℝ^*c×H×W*^, where *I* = *I*_st_ when only transcriptomic data are used (*c* = *p* + 1), and *I* = (*I*_st_, *I*_histo_)_channel_ when histology is included (*c* = *p* + 2), with concatenation performed along the channel dimension. These unified rasters provide a continuous, resolution-adaptive representation on which all subsequent optimisation is performed.

### 2.3. Alignment objective and optimisation

GALA models the spatial correspondence between source and target raster tensors through a composite transformation (*ϕ*^*v*^ *· A*)(***x***) = *ϕ*^*v*^(*A*(***x***)). The transformation consists of a global affine matrix *A* to correct large-scale differences (rotation, translation, scaling, and reflection) with a diffeomorphic deformation *ϕ*^*v*^ parameterised by a time-varying velocity field *v*_*t*_ (*t* ∈ [0, 1]) for each spatial location ***x*** ∈ ℝ^2^ (see Fig. 1b and Supplementary Section S1.2).

Building on the global affine alignment that estimates the overlapping region between source and target, GALA restricts subsequent refinement to this overlap domain. Within the estimated overlapping region, a probabilistic matching model is introduced to distinguish matched (reliably aligned) areas from unmatched (background, missing, or artefactual) areas. Specifically, for each grid location ***x*** ∈ Ω, a latent label indicates whether it belongs to the matched or unmatched class. Let *I*(***x***), *J*(***x***) ∈ ℝ^*c*^ denote the raster intensities of source and target at location ***x***. Under the composite transformation (*ϕ*^*v*^ *·A*)(***x***), we model the observed target intensity as

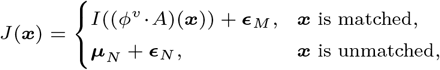

where the residuals ***ϵ*** are defined as deviations from their respective mean intensities. To quantify similarity, we evaluate these residuals using Gaussian kernels 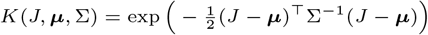 with mean ***µ*** and covariance ∑. ***µ***_*N*_ denotes the mean intensity of unmatched areas, estimated iteratively. This Gaussian form induces a quadratic data term and enables closed-form Expectation-Maximisation (EM) updates [12, 35]. Therefore, given class priors *π*_*M*_ and *π*_*N*_ (*π*_*M*_ + *π*_*N*_ = 1), the posterior probability that location ***x*** belongs to the matched class is computed as

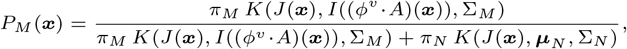

where 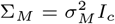 and 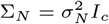 are the covariance matrices of ***ϵ***_*M*_ and ***ϵ***_*N*_, respectively, *I*_*c*_ is the *c × c* identity matrix. A high value of *P*_*M*_ (***x***) indicates strong correspondence, whereas a low value suggests background or artefactual areas.

To incorporate the probabilistic matching model into the alignment procedure, the posterior probability *P*_*M*_ (***x***) is used to weight the contribution of each spatial location in the optimisation. The multimodal alignment objective combines transcriptomic and histological fidelity terms (weighted by *α* and 1 − *α*) with a regularisation on the deformation field *v*:

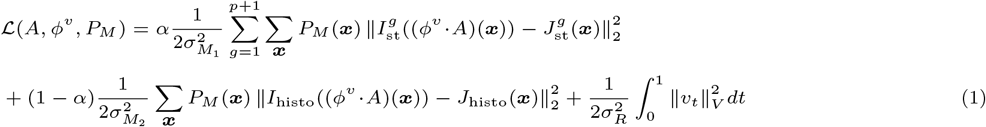

where 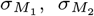, and *σ*_*R*_ control feature matching and deformation smoothness. The first two terms minimise transcriptomic and morphological discrepancies between the iteratively aligned source and target, weighted by the matching probability *P*_*M*_ (***x***). The final term regularises *v*_*t*_ in a reproducing kernel Hilbert space (RKHS), ensuring smooth and invertible mappings [36]. The selected *p* + 1 gene channels and histological image channel act as alignment “landmarks”, guiding the optimisation—principally differing from STalign, which does not utilise this information. Additional details are provided in Supplementary Section S1.2.

We adopt a coupled optimisation strategy that jointly refines global affine alignment and local diffeomorphic deformation within a unified objective. In the global stage, a genetic algorithm (GA) [37] conducts a population-based search to estimate affine parameters and overlapping regions, providing robust initialisation that avoids local minima. Subsequently, the local stage resolves residual nonlinear discrepancies using large deformation diffeomorphic metric mapping (LDDMM) [36], modelling smooth, invertible deformations suitable for fine-grained spatial adjustments. To reconcile local deformations with modality-specific information, an EM-like LDDMM scheme is implemented: in the E-step, matching probabilities *P*_*M*_ are updated from current residual kernels; in the M-step, the velocity field *v*_*t*_ is optimised by minimising the weighted quadratic registration energy under the LDDMM framework, while ***µ***_*N*_ is updated accordingly. The pseudo-code of GALA is provided in Supplementary Alg. S1.

Unlike conventional pipelines that perform global (coarse) and local (fine) alignment independently, our coordinated coarse-to-fine scheme iteratively reinforces both stages. This bidirectional coupling, where coarse estimates guide local refinements and local updates feedback to adjust global parameters, enables more stable convergence than purely hierarchical or decoupled approaches. A full mathematical formulation and optimisation procedure are provided in Supplementary Section 1.3. Empirically, under default parameter settings (Supplementary Tab. S1), only two global–local episodes suffice to achieve stable alignment across most datasets (Supplementary Fig. S3).

### 2.4. Selection of informative genes and hyper-parameters

GALA requires two key hyper-parameters: the number of selected informative genes *p*, which guide alignment, and the relative weight *α* that balances contributions from transcriptomics profiles versus histological images.

For gene selection, we adopted the *R*^2^-based strategy from GPSA [21], which ranks genes by their predictability from local spatial context. Specifically, for each gene, a *k*-nearest neighbour (*k*NN) regression model is trained to predict expression levels from the spatial coordinates of neighbouring spots or cells. The coefficient of determination (*R*^2^) quantifies spatial predictability, and the top *p* genes with the highest *R*^2^ values from the source data are retained. Alternative approaches, such as identifying spatially variable genes (SVG) [38, 39] or using known marker genes, are also viable; we chose the *R*^2^-based method for its simplicity, computational efficiency, and robust performance across diverse datasets, as previously validated by GPSA. Systematic sensitivity analysis indicated that *p* = 3 (top three informative genes) suffices even under partial alignment scenarios, and this value is set as default. Notably, GALA does not require these informative genes to provide full anatomical coverage or be spatially complementary, as the inclusion of a total expression channel provides a consistent global structural scaffold. The balance parameter *α* was observed to yield stable performance across its range (0, 1); we set *α* = 0.5 when histological image data are available, and *α* = 1 otherwise. Detailed robustness and sensitivity analyses, as well as spatial coverage analysis of informative genes, are provided in Supplementary Sections S2.1.1, S2.1.2, and S2.1.4.

### 2.5. Evaluation metrics

To evaluate alignment performance across diverse experimental scenarios, we employ complementary metrics capturing spatial accuracy, biological coherence, and molecular consistency. Metrics are selected based on dataset characteristics, such as availability of anatomical annotations, spatial resolution, and modality, to ensure biologically meaningful and context-appropriate assessment.

When anatomical labels or ground truth are available, we assess alignment fidelity using two strategies. First, label consistency accuracy identifies mutual nearest-neighbour pairs between aligned slices and reports the proportion sharing the same anatomical label, directly quantifying structural correspondence and, more specifically, the preservation of biologically defined regions [8]. Second, biological coherence is evaluated via clustering-based metrics: a joint Gaussian mixture module (GMM) is fitted to combined spatial and gene expression features from both aligned source and target slices, and cluster assignments are compared with ground-truth labels using Adjusted Rand Index (ARI) and Normalized Mutual Information (NMI). These metrics assess how well alignment preserves region-specific transcriptional and anatomical structure.

For spot-based datasets lacking reliable anatomical annotations, spatial cross-correlation metrics quantify preservation of continuous spatial gene expression gradients [40, 41]. For each gene, correlations between transcriptomic profiles in aligned source and target slices are computed with Gaussian-kernel weights based on inter-spot distances. Averaging across genes provides a global measure of spatial expression consistency, capturing local gradient preservation without requiring labels. Further details on evaluation metrics are provided in Supplementary Section S1.4.

In cell-based datasets, direct cell-to-cell comparison is often unreliable due to biological heterogeneity and measurement sparsity. Following STalign[12], neighbouring cells are aggregated into pseudo-spots to stabilise expression signals. Cosine similarity between corresponding pseudo-spots across aligned slices then quantifies local molecular agreement, mitigating single-cell noise and providing an effective fidelity measure for high-resolution single-cell datasets without histological or anatomical ground truth.

For transcriptomics-to-histology alignment, we use the Mean Absolute Error (MAE) of annotated landmark coordinates:

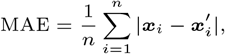

where *i* = 1, …, *n* indexes landmarks, and 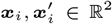 denote the coordinates of the *i*-th landmark in the target and aligned source data, respectively. Lower MAE indicates better spatial correspondence and provides an evaluation that is independent of gene expression or clustering assumptions, making it compatible with landmark-free alignment methods.

Overall, each metric is selected to match the characteristics of the corresponding dataset and alignment task, capturing structural correspondence (label consistency, MAE), transcriptional coherence (ARI, NMI, pseudo-spot similarity), or preservation of spatial expression gradients (cross-correlation). These quantitative measures are complemented by visual overlays of aligned slices, providing intuitive verification of spot/cell correspondence, gene expression patterns, and regional morphology. Together, they offer a comprehensive assessment of GALA’s performance across diverse spatial transcriptomics datasets.

### 2.6. Implementation techniques for all alignment scenarios

GALA is a versatile alignment framework applicable to widespread application scenarios (see Fig. 1c). For example, whether the source and target data i) belong to the same modality or not, ii) are consistent in spatial resolution or not, and iii) are fully or partially overlapped. In the following, we detail how GALA is applied across these scenarios.

First, because rasterisation maps each dataset onto a user-defined grid, same-modality alignment under both matched and mismatched resolutions, including spot-to-spot, cell-to-cell, and cell-to-spot scenarios, is handled directly through the choice of grid spacing *dx* (see Section 2.2). No additional algorithmic modifications are required: once rasterised, all datasets are processed identically by the optimisation pipeline. This design enables resolution-aware and platform-agnostic alignment across Visium, Xenium, MERFISH, and related technologies. Second, GALA supports cross-modality alignment, such as registering spatial transcriptomic measurements to histological images from the same tissue section. Histology images are converted to greyscale and transformed into structural contrast maps, whereas transcriptomic measurements are encoded as rasterisation tensors, allowing both to be treated as image-like modalities. Alignment is then performed under the multimodal objective:

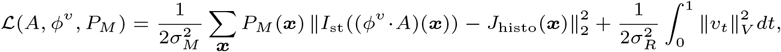

where 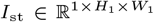 and 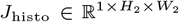. This variant enables cross-modality alignment without requiring landmarks or manual correspondence, facilitating integration across different types of measurements in ST studies.

Third, GALA explicitly accommodates partial alignment. The framework first performs a population-based affine parameter search to approximate the overlapping region between the two datasets. This global affine estimate then guides the subsequent LDDMM-based refinement. During refinement, an EM-like procedure updates pixel-wise correspondence weights, allowing the model to concentrate on structurally matched areas while downweighting unmatched or noisy areas (see Section 2.3). This mechanism confers robustness in scenarios involving incomplete, distorted, or physically damaged tissue.

Finally, GALA’s multimodal rasterisation and flexible objective function allow natural extension to other omics modalities beyond transcriptomics. For instance, metabolic or proteomic imaging data can be rasterised similarly, enabling alignment of metabolite distributions or protein expression patterns to transcriptomic or histological references. This generalisability makes GALA a broadly applicable framework for multi-omics spatial alignment, supporting integrative studies across diverse molecular modalities and imaging platforms.

## 3. Results

We systematically evaluate GALA across a wide range of ST alignment tasks (Fig. 1c) and compare its performance with state-of-the-art methods using established quantitative metrics (Section 2.5). Because existing alignment methods are typically designed for specific settings, the set of baseline methods differs across alignment scenarios.

### 3.1. Spot-to-spot alignment within ST technologies

#### 3.1.1. Human dorsolateral pre-frontal cortex (DLPFC) data by Visium

The first example used to evaluate GALA is the human dorsolateral pre-frontal cortex (DLPFC) ST data generated with the 10x Genomics Visium platform [29]. The dataset comprises three independent samples (A, B, and C), each containing four consecutive tissue slices annotated into seven anatomical regions: six neocortical layers and white matter (WM). Each slice contains 3,000–4,000 spots and approximately 10,000 genes. The spacing between adjacent slices is 10 *µ*m for pairs 1–2 and 3–4, but increases to 300 *µ*m between slices 2 and 3, introducing both gradual and abrupt structural variation (see Sample C in Fig. 2a). We focus on Sample C due to its complete annotation, clear laminar structure, and widespread use in recent comparative studies [13, 20, 42, 43]. GALA was compared against five baseline methods applicable to spot-to-spot alignment, spanning rigid (PASTE, STAligner), non-rigid (GPSA), one-to-one matching (SLAT), and coarse-to-fine (ST-GEARS) strategies. Evaluation metrics include label consistency accuracy, ARI, and NMI to quantitatively assess biological coherence after alignment (see Section 2.5).

**Fig. 2.**
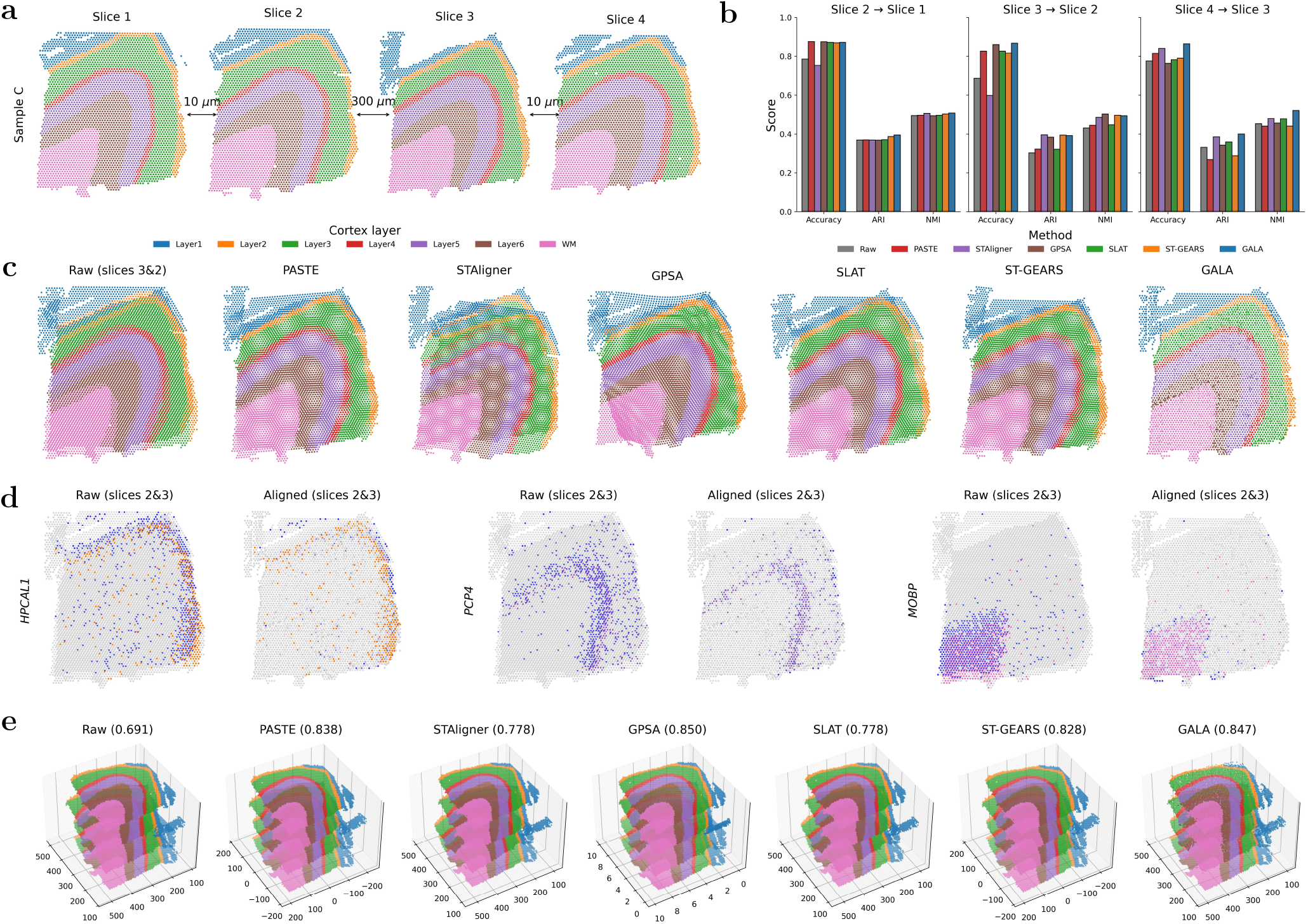
Alignment evaluation on Sample C of DLPFC data across GALA and baselines. **a** Four raw tissue slices from Sample C, with adjacent distances of 10 *µ*m, 300 *µ*m, and 10 *µ*m. Spots are annotated with six neocortical layers (Layer 1–6) and white matter (WM). **b** Quantitative comparison of alignment performance for three pairwise slice alignments (2 → 1, 3 → 2, 4 → 3) based on label consistency accuracy (Accuracy), ARI, and NMI. ‘Raw’ indicates unaligned slices. **c** Visual comparison of aligning slice 3 to slice 2 across all methods (except ‘Raw’). The differently sized white background dots result from the stacked two regular grids following affine transformation, generating a Moiré-like pattern. **d** Visual comparison for spatial distribution of the three layer marker genes before and after alignment by GALA. Spots with high marker-gene expression in the target slice are blue; low-expression spots are grey. In the source and aligned source slices, high-expression spots are coloured according to their annotated layer in **a**, while low-expression spots are grey. **e** 3D visualization of all four slices reconstructed via sequential pairwise alignment using GALA and baselines. The accuracy values shown in parentheses after each method denote the mean pairwise label consistency accuracy.

We first considered three pairwise slice alignments (2 → 1, 3 → 2, and 4 → 3) for Sample C without histological images. As shown in Fig. 2b, GALA achieves the highest mean label consistency accuracy (0.868) across these comparisons, surpassing top-performing methods such as PASTE (0.839) and GPSA (0.833). Clustering consistency metrics further supported GALA’s advantage, with joint GMM analysis yielding a mean ARI of 0.396 and NMI of 0.508, outperforming STAligner (0.385 and 0.491, respectively; see Supplementary Tab. S5 and Fig. S10a for detailed metric values). These results demonstrate that GALA preserves label consistency and biological coherence more effectively than all baselines. Visual comparison of aligning slice 3 to slice 2 (Fig. 2c) shows that GALA achieves the most precise alignment of all major cortical layers, whereas STAligner exhibits global structural mismatches and GPSA produces substantial distortions in WM (pink). Visual comparisons for slice pairs 2 → 1 and 4 → 3 are provided in Supplementary Fig. S10b-c.

We further assessed biological consistency by examining canonical layer-specific marker genes. As shown in Fig. 2d, stacked source and target slices display markedly improved spatial correspondence after GALA alignment. Spots with high marker-gene expression are shown in blue on the target slice and are coloured by their annotated layer on the source and aligned source slices; low-expression spots are shown in grey in all panels. Representative markers (*HPCAL1, PCP4, MOBP*) exhibit increased overlap after alignment, indicating enhanced layer preservation and improved biological coherence. Using pairwise alignment, 3D visualization was performed by setting slice 1 as the template and aligning other slices to a common coordinate system, shown in Fig. 2e. Rather than a dedicated 3D reconstruction method, GALA achieves 3D structural consistency that is comparable to specialised 3D integration approaches, including STaligner, GPSA, and ST-GEARS. In particular, cross-slice layer transitions and anatomical boundaries remain well preserved across adjacent sections. Quantitative pairwise label consistency results further support this observation, showing that GALA (0.847) performs competitively and differs only marginally from GPSA (0.850), consistent with the subtle visual differences observed in the 3D rendering.

Next, we evaluated the effect of incorporating H&E-stained histology images on alignment between slices 3 and 2. GPSA was included as the only baseline supporting image integration. As shown in Fig. 3a (bar chart), GALA achieved label accuracies of 0.869 and 0.875 with and without histology, respectively, outperforming GPSA (0.833 and 0.830). Visual inspection confirms that GALA produces more accurate and smoother alignments, indicating that inclusion of morphological information benefits GALA while introducing greater variability in GPSA’s Gaussian process. As discussed in Section 2.4, the fusion weight *α* is set to 0.5 by default when histology is available. Additional experiments (line chart in Fig. 3a) show GALA remains stable for *α* near 0.5, demonstrating robustness to the relative weight.

**Fig. 3.**
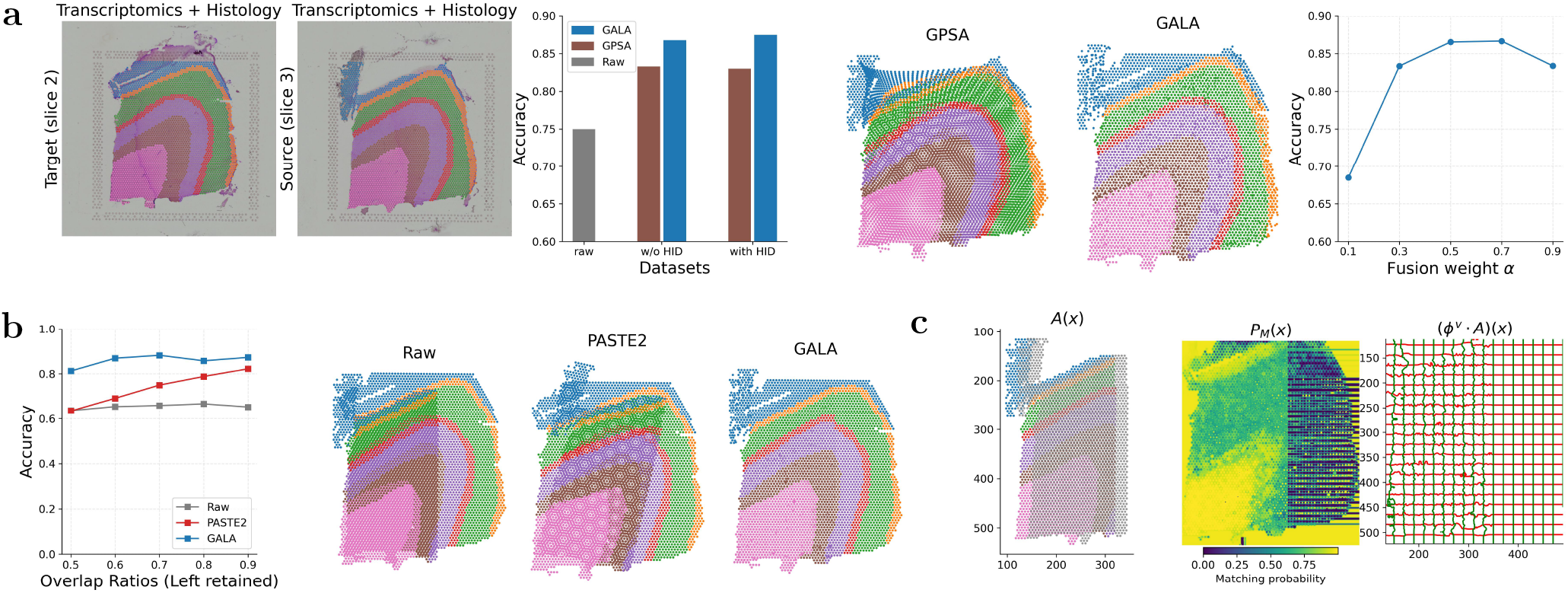
Multimodal alignment and robustness to partial overlap using GALA. Spots are coloured according to their annotated layer in Fig. 2**a**. **a** Comparison of alignment between slices 3 and 2 when histological image data are included, across GPSA and GALA. The rightmost panel shows sensitivity to the relative weight *α*. ‘w/o HID’ indicates no histology. **b** Label consistency accuracy for partial alignment (left-retained slice 3) across overlap ratios 50%–90%; right panels show the visual comparison of the corresponding partial alignment in the case of overlap ratio 60%. **c** Illustration of the GALA mechanism for ‘left 60% retained’: global affine transformation *A*(***x***), matching probabilities *P*_*M*_ (***x***), and local diffeomorphic mapping (*ϕ*^*v*^ *·A*)(***x***).

Third, we explored partial alignment by cropping slice 3 to retain either the left or bottom regions across overlap ratios of 50–90%, incorporating histology in both source and target. GALA was compared with PASTE2, a recent method designed for partial ST alignment. Fig. 3b shows label consistency accuracy for raw data, PASTE2, and GALA when the left section is retained; results for the bottom section are in Supplementary Fig. S11a. Across all overlap ratios in both partial settings, GALA consistently achieved higher and more stable accuracy than PASTE2. Visual results for 60% overlap further demonstrate that only GALA preserves laminar continuity with minimal distortion. Sensitivity analysis of *α* (Supplementary Fig. S11c) confirms robustness consistent with Fig. 3a.

Finally, we illustrated why GALA performs well in partial alignment by visualising learned parameters for a 60% overlap (first type of partial overlap; second type shown in Supplementary Fig. S7b). The left panel of Fig. 3c and Supplementary Fig. S11b shows the global affine transformation of the cropped source from the original location (grey) to the transformed location (coloured) in the target coordinate system. The middle panel indicates the learned matching probability *P*_*M*_ for each target spot, with higher values (brighter) highlighting the reliably matched areas. The right panel shows the local refinement via the learned diffeomorphic mapping (*ϕ*^*v*^ *· A*), producing smooth, topology-preserving deformations. Only the lattice in the overlapped part, corresponding to high *P*_*M*_ values, is distorted. The global affine transformation and local diffeomorphic mapping are iteratively optimised under the objective function, explaining GALA’s effectiveness for partial ST alignment.

#### 3.1.2. Mouse brain sagittal-posterior (MBSP) sections by Visium

The second example of spot-to-spot alignment involves mouse brain sagittal-posterior (MBSP) sections produced by 10x Genomics Visium. The two sections contain 2,805 and 2,840 spots, respectively, each profiled for 32,285 genes, and are accompanied by H&E-stained histological images (left panel of Fig. 4a).

**Fig. 4.**
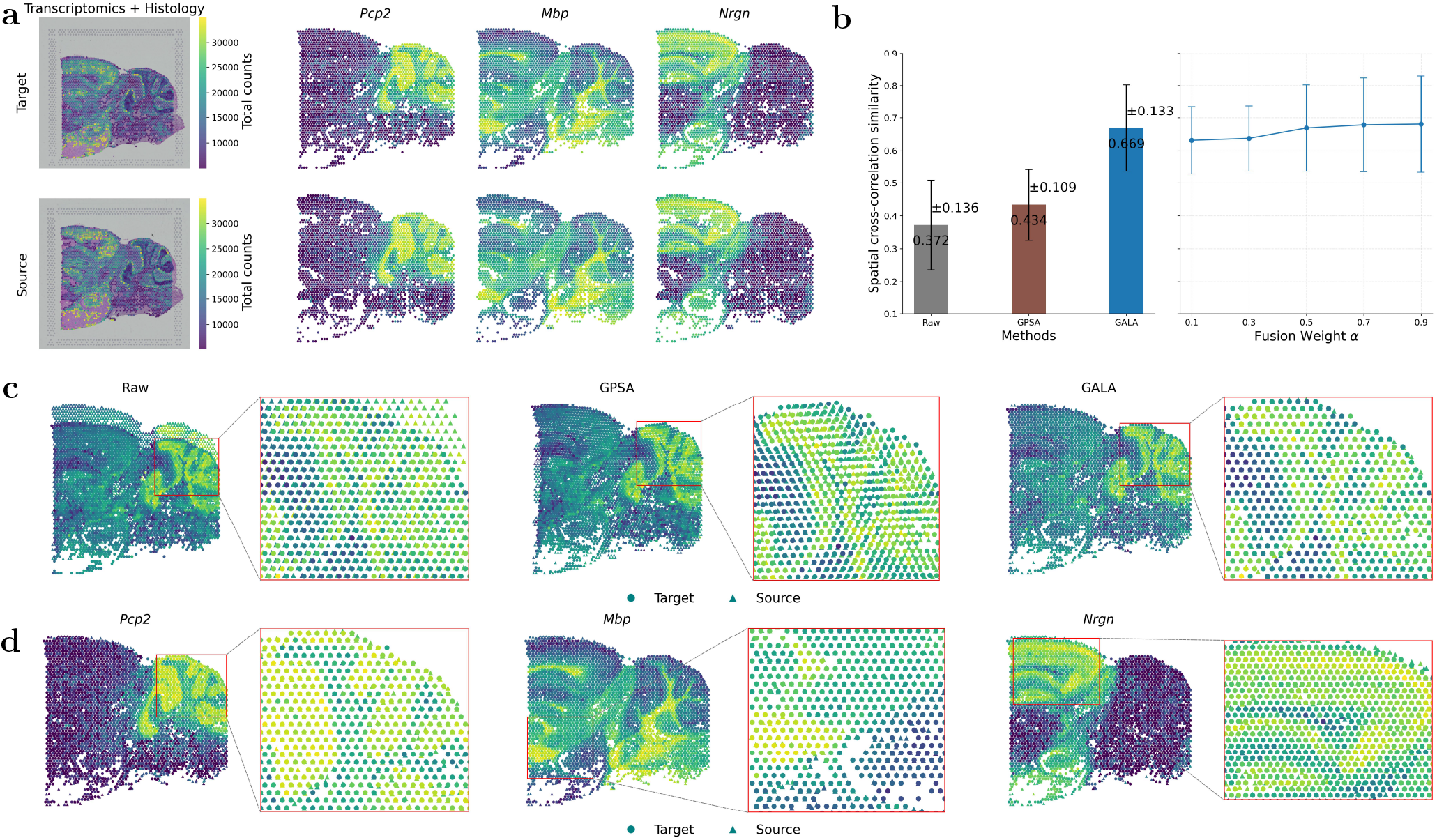
Alignment of two MBSP sections. **a** Raw slices overlaid on H&E images and spatial expression of three top informative genes (*Pcp2, Mbp, Nrgn*). Brighter spots indicate higher gene expression. **b** Left: mean spatial cross-similarity scores (± s.d.) across the top 135 informative genes for raw slices (Raw) and slices aligned by GPSA and GALA (*α* = 0.5). Right: cross-similarity ± s.d. across fusion weights from 0.1 to 0.9. **c** Visual comparison of alignment: left, raw stacked slices; middle and right, slices aligned by GPSA and GALA. Zoomed regions correspond to red boxes. **d** Visual alignment of the three informative genes by GALA, with zoomed-in regions for each panel.

We first examined the role of spatially informative genes in GALA, defined as genes ranked by the *R*^2^-based strategy from GPSA [21], which orders genes according to how well their expression can be predicted from local spatial context (see Section 2.4). The top three genes automatically selected were *Pcp2, Mbp*, and *Nrgn*. These genes exhibit complementary anatomical specificity (Fig. 4a, right panels): *Pcp2* is restricted to cerebellar Purkinje cells and selected cortical layers; *Mbp* marks oligodendrocyte-enriched white matter tracts; *Nrgn* is expressed in excitatory neurons within cortical and hippocampal layers. Their distinct localisation patterns provide strong signals for guiding alignment, effectively serving as ‘landmarks’ in our landmark-free GALA framework.

Unlike the previous example, ground-truth spot annotations of MBSP sections are unavailable. Therefore, we used spatial cross-correlation similarity (instead of label consistency accuracy, ARI, or NMI) to quantify alignment quality (see Section 2.5). Using GPSA’s protocol, we selected the top 135 informative genes with *R*^2^ *>* 0.6, each reporting one correlation score after alignment. We compared GALA with GPSA only, as no other baselines support histological image data. The bar charts in Fig. 4b show that: (i) both methods substantially improve alignment relative to raw slices (mean 0.372 ± 0.136); (ii) GALA achieves a higher mean score of 0.669 ± 0.133, an improvement of over 23% relative to GPSA (0.434 ± 0.109). Visual comparison of stacked target and aligned source slices (Fig. 4c) highlights that anatomical features are highly displaced in raw slices. GPSA corrects misalignment globally but shows local distortions, whereas GALA provides finer alignment with better preservation of structural boundaries.

We next examined the alignment of the top three informative genes (Fig.4d). Compared with their raw, unaligned spatial patterns (see Supplementary Fig. S12), all three genes display markedly improved localisation following GALA alignment. In the cerebellar cortex, *Pcp2* becomes more sharply confined to Purkinje cell layers, with clearer lobular boundaries and reduced cross-layer bleeding. *Mbp*, which marks white-matter tracts, shows more continuous and anatomically coherent fibre bundles, while *Nrgn* exhibits a smoother and more spatially consistent distribution across granule and cortical layers. These improvements, further highlighted in the magnified insets (red boxes), demonstrate GALA’s ability to recover biologically coherent expression domains by correcting structural variability across slices. Sensitivity analysis of the fusion weight *α* (default 0.5) further confirmed GALA’s robustness, as shown in Fig.4b.

### 3.2. Cell-to-cell alignment within ST technologies

In the third example, we applied GALA to subcellular-resolved ST data from the MERFISH mouse liver map datasets (Vizgen) without histological image data. The two liver slices, L2S1 and L2S2 from Animal 2, each comprises approximately 300,000 cells with expression profiles for 347 genes, serving as target and source data, respectively. Fig. 5a shows the raw slices, displaying only the positions of cells with detected gene expression (the corresponding total gene expression maps are provided in Supplementary Fig. S13a), highlighting substantial rotation. Baseline methods include SLAT, STalign, and the affine component of GALA (denoted as Affine).

**Fig. 5.**
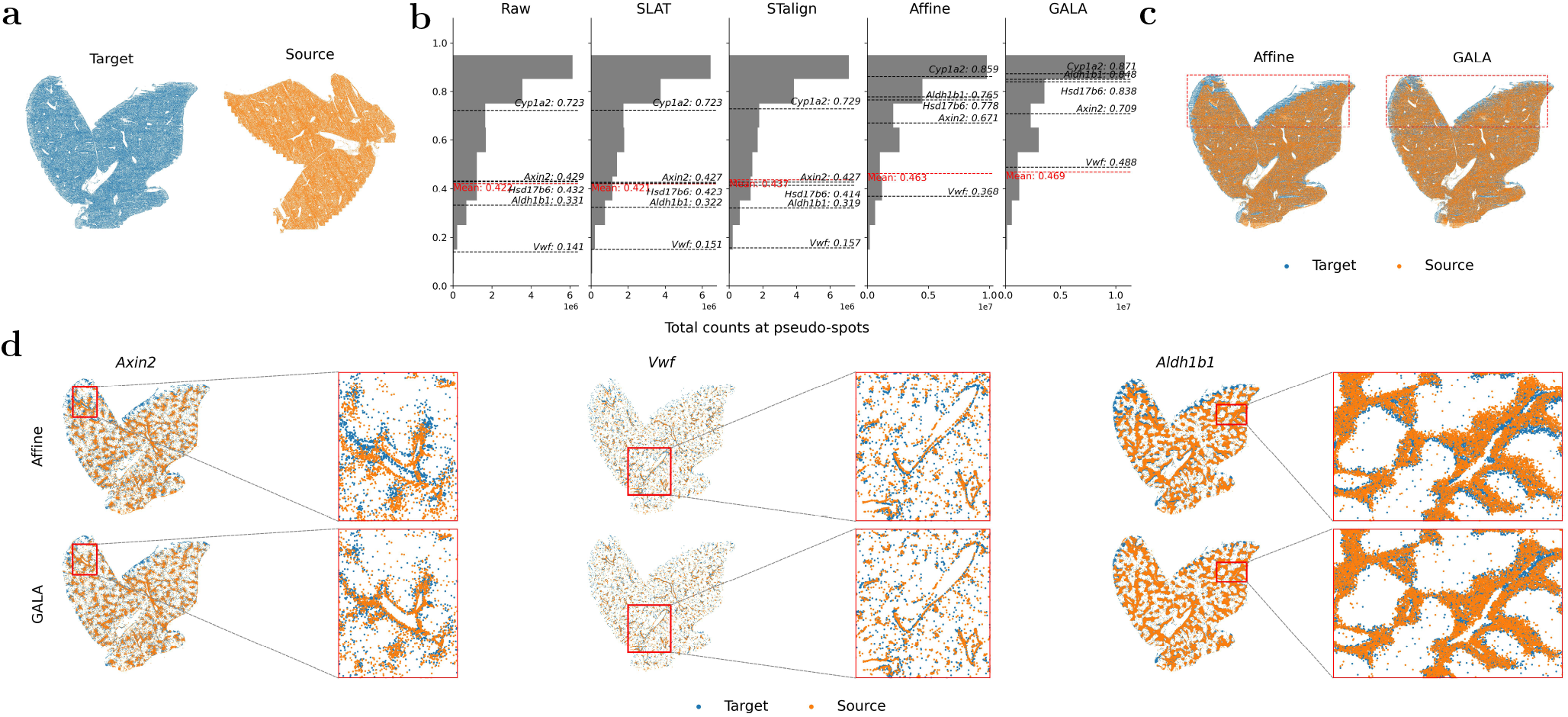
Alignment of two MERFISH mouse liver replicates. **a** Raw slices for source and target data, coloured in blue and orange, respectively. Scaled expression slices are shown in Supplementary Fig. 13a. **b** Distribution of cosine similarity scores for 134 liver marker genes and 3 informative genes against the total gene expression at pseudo-spots (aggregated into 30 *µ*m) after alignment by Raw, SLAT, STalign, Affine, and GALA. Red and black dashed lines indicate the mean and representative genes, respectively. **c** Visual comparison of whole-slice alignment results by Affine and GALA. Cells in target and aligned source data are indicated as blue and orange, respectively. The top area in red rectangular highlights the significant improvement of local transformation from GALA. **d** Alignment of three representative genes at whole-slice and zoomed-in regions, comparing Affine and GALA.

To quantify alignment quality, cells were aggregated into uniform 30 *µ*m pseudo-spots to mitigate local sampling density variation, following STalign’s protocol. Cosine similarity of expression was computed between matched pseudo-spots, capturing local correspondence in both gene expression magnitude and spatial structure. Evaluation considered 134 known liver marker genes [44] and top-ranked informative genes. Five representative genes, including canonical markers (*Axin2* [45], *Vwf* [46]) and top informative genes (*Aldh1b1, Hsd17b6, Cpy12a*), were analysed in detail.

First, we quantitatively assessed alignment across all baselines and GALA. Fig. 5b shows the distribution of cosine similarity scores for 134 liver marker genes against the total gene expression at pseudo-spots after alignment by these methods (including Raw data). The mean similarity score and the scores for the five representative genes are indicated by red dashed line and black dashed lines, respectively. SLAT yielded negligible improvement: its one-to-one matching is sensitive to variations in cell density and can introduce spurious correspondences. STalign delivered only marginal gains because it depends on manually specified priors to guide deformation, which limits performance when prior correspondence is imprecise. The affine-only variant of GALA (Affine) modestly increased the mean similarity to 0.463, and the full GALA framework further raised it to 0.469, demonstrating that both global affine and local diffeomorphic components contribute to alignment. Representative gene trends exemplify this: raw alignment produced scores of 0.429 (*Axin2*), 0.141 (*Vwf*), 0.331 (*Aldh1b1*), 0.432 (*Hsd17b6*), and 0.723 (*Cyp12a*); Affine improved these to 0.671, 0.368, 0.765, 0.778, and 0.859; full GALA further increased them to 0.709, 0.488, 0.848, 0.838, and 0.871.

Second, we visually compared alignment results for Affine and GALA. The right panels of Fig. 5c show that both methods achieve global alignment, but GALA produces better local alignment, with the top region (red box) showing more coherent overlap and reduced spatial drift. Fig. 5d illustrates alignment for three representative genes, including zoomed-in regions, showing that GALA more accurately aligns source and target cells for canonical markers *Axin2, Vwf* and informative gene *Aldh1b1*. (Supplementary Fig. S13b, c illustrate the corresponding gene expression values).

These results demonstrate that GALA’s coupled coarse-to-fine optimisation improves global alignment accuracy while preserving local gene-level patterns in highly resolved single-cell spatial transcriptomic datasets, even without H&E guidance or manual landmarks.

### 3.3. Cell-to-spot alignment across ST technologies

GALA also supports mismatched resolution alignment across technologies, i.e., aligning single-cell resolved ST data (e.g. from MERFISH, Xenium) to multi-cellular resolved ST data (e.g. from Visium). Such alignment enables mapping of cell types and gene expression domains across resolution scales, supporting comparative studies between experiments and modalities [20, 24, 47]. In the fourth example, the source and target data are Fresh Frozen mouse brain replicate 2 from the Xenium platform and a mouse coronal section from the Visium platform, respectively. The Xenium dataset contains 149,292 single cells with high-resolution gene expression, whereas the Visium data comprise 2,634 spots with low-resolution expression profiles. As shown in Fig. 6a, the Xenium slice spans a full coronal brain section, while the Visium slice captures only cortical and subcortical structures in the left hemisphere, making this a partial alignment across resolutions. Baseline methods include SLAT and Affine. STalign was not considered because it cannot directly perform cell-to-spot alignment; instead, it aligns single-cell ST data to the registered H&E image corresponding to the multi-cellular ST data, requiring manually selected landmarks.

**Fig. 6.**
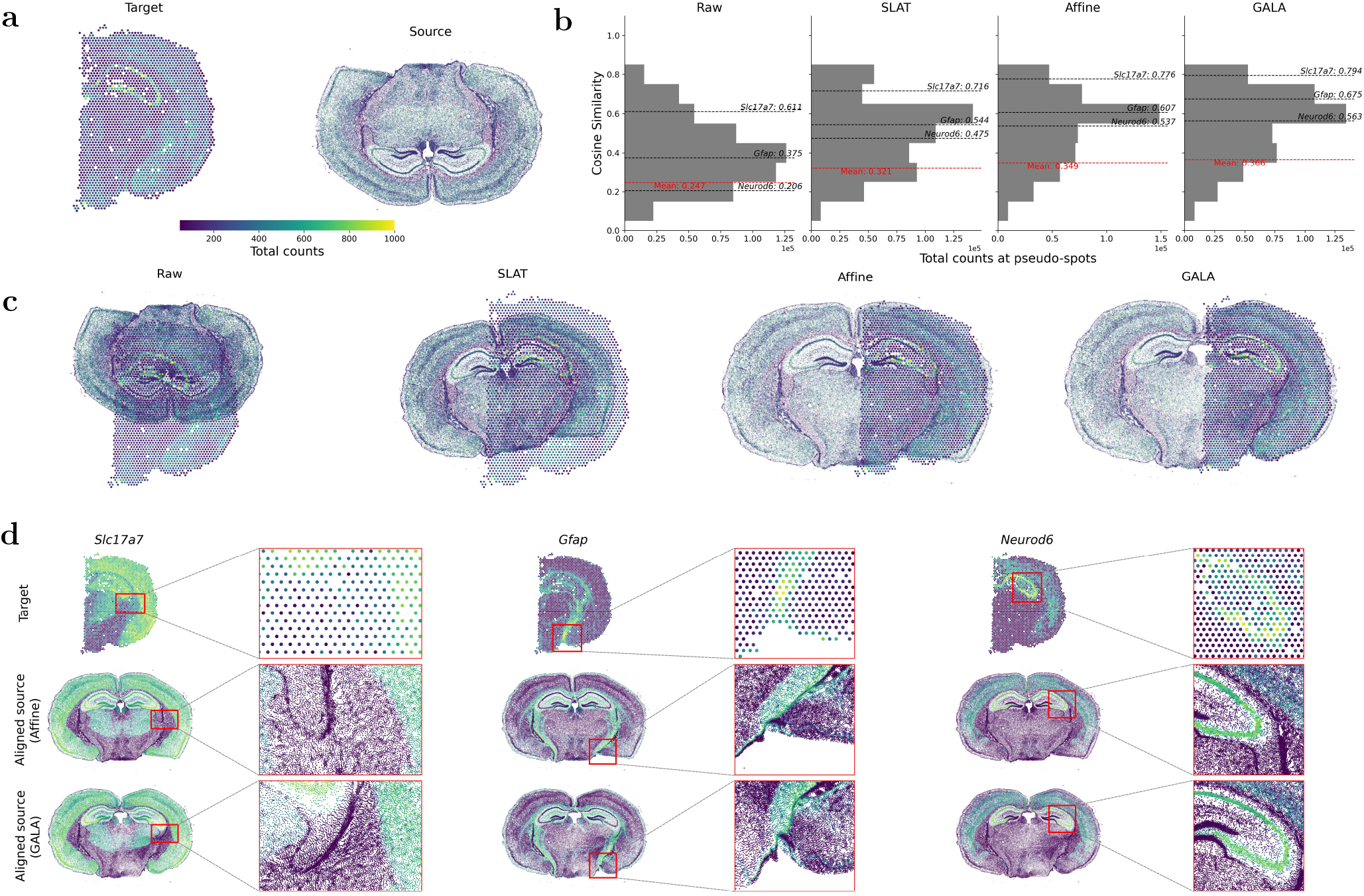
Alignment of mouse coronal sections from Visium and Xenium. **a** Visualisation of Visium (spot-level, target) and Xenium (cell-level, source) slices. Brighter colours indicate higher gene expression. **b** Distribution of cosine similarities between gene expression in Visium target versus aligned Xenium source at matched spots/pseudo-spots. Red dashed lines indicate mean similarity; black dashed lines highlight three informative genes. **c** Comparison of unaligned slices with slices aligned by SLAT, Affine, and GALA. **d** Spatial expression of three informative genes, *Slc17a7, Gfap*, and *Neurod6*, in Visium (top), Affine-aligned (middle), and GALA-aligned Xenium data (bottom), with magnified regions indicated by red rectangles.

To quantify alignment, we computed cosine similarity of gene expression vectors aggregated into 30 *µ*m pseudo-spots for all common genes (Fig. 6b). Mean similarity scores show that GALA and Affine substantially outperform Raw and SLAT. We also examined alignment of three spatially informative marker genes: *Slc17a7* (excitatory cortical neurons), *Gfap* (astrocytes near the ventricular zone and midline), and *Neurod6* (hippocampus and olfactory bulb). Fig. 6b demonstrates that, compared with raw similarities of 0.611, 0.375, and 0.206, GALA increased these to 0.794, 0.675, and 0.563, outperforming Affine-aligned results of 0.776, 0.607, and 0.537, and SLAT-aligned results of 0.716, 0.544, and 0.475.

We further visually evaluated alignment across methods. As shown in Fig. 6c, SLAT performs poorly in this cross-resolution setting, primarily due to its reliance on one-to-one adversarial graph matching followed by a global affine transformation, which introduces noise and fails to capture local nonlinear deformations. In contrast, GALA and Affine achieve markedly improved global alignment, with GALA producing finer geometric correspondence and better preservation of fine-scale structural coherence. Supplementary Fig. S14 provides additional comparisons, showing target data as registered H&E images. Fig. 6d demonstrates alignment at the molecular level for GALA and Affine. Regions highlighted by red boxes exemplify alignment differences, with GALA achieving more coherent correspondence between aligned source and target data.

These results indicate that in the cell-to-spot alignment scenario, GALA outperforms baseline methods, and both components of the global affine and local diffeomorphic refinement contribute substantially to the improved alignment. These conclusions are consistent with observations in the cell-to-cell scenario in Section 3.2.

### 3.4. Cell-level transcriptomics to histological image alignment within the same tissue section

Lastly, we evaluated GALA for aligning ST data to paired H&E images within the same tissue section and experimental platform. Such registration is necessary to correct physical and technical misalignments arising from the automated instrument workflow, rather than biological variation [12, 28]. Misalignments are primarily rigid-body transformations (translation and rotation) introduced during mechanical handling, with minor scaling effects possible due to optical calibration. Accurate alignment under this scenario bridges the gap between the abstract molecular map and interpretable tissue morphology.

We applied GALA to two datasets: an FFPE human lung cancer section (Experiment 1, Xenium In Situ) and an FFPE human breast cancer section (Replicate 1, Xenium In Situ Sample 1). Fig. 7a and c show the H&E image (left; target) and the corresponding high-resolution single-cell expression data (right; source). Clear differences in scale, rotation, and translation are visible, while Fig. 7c also shows a reflection between the expression and histology coordinates, likely arising from slide orientation or scanning artefacts.

**Fig. 7.**
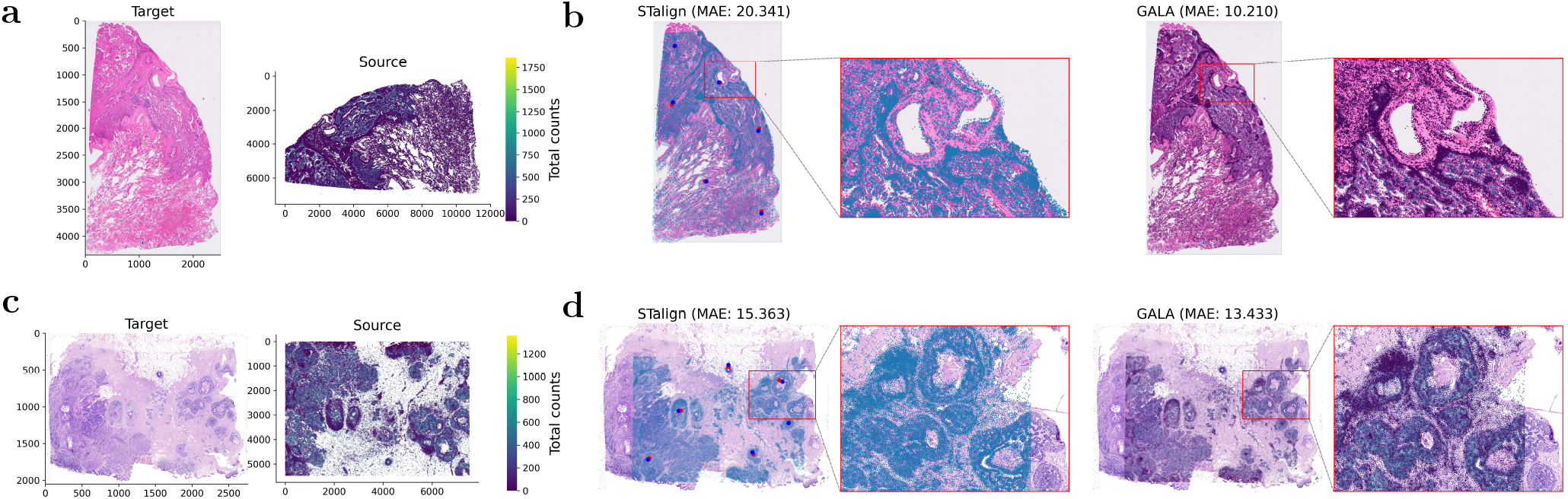
Alignment of Xenium single-cell RNA data to paired H&E images. **a** H&E histological image (target) and paired Xenium single-cell data (source) from a human lung cancer section, where brighter colours indicate higher total gene expression. **b** Alignment results of **a** by landmark-based STalign and landmark-free GALA, with MAE indicated. Blue/red dots in the STalign panel denote manually annotated landmarks in source/target data. As STalign ignores gene expression, its aligned data are visualised as a blue overlay on the target H&E image. Adjacent zoom-in panels highlight regions marked by red boxes. **c** H&E image (target) and paired single-cell data (source) from a human breast cancer section, with brighter colours indicating higher expression. **d** Alignment results of **c** by STalign and GALA, with MAE indicated and magnified regions within red rectangles.

To our knowledge, besides GALA, only STalign addresses the problem of transcriptomics-to-histology alignment. However, STalign requires manually annotated landmarks for affine transformation without utilizing gene expression, whereas GALA is fully landmark-free and incorporates gene expression. We compared STalign and GALA using the MAE of landmark coordinates to quantitatively assess alignment (see Section 2.5). Fig. 7b shows the lung cancer dataset results: using six representative landmarks (compared with the four landmarks used for pre-alignment in STalign), STalign achieved an MAE of 20.341, whereas GALA achieved a more precise alignment with MAE of 10.210. Fig. 7d further demonstrates GALA’s superior performance on the breast cancer dataset, with MAEs of 15.363 and 13.433 for STalign and GALA, respectively.

Landmark selection is inherently subjective and prone to user bias. Different experts may select different landmarks, leading to inconsistent alignments. For instance, when landmarks are sparse or lack anatomical specificity, STalign’s performance degrades markedly (Supplementary Fig. S15). In contrast, GALA does not rely on manual landmarks during optimisation, eliminating user bias and ensuring fully data-driven alignment. Landmarks are used only for independent evaluation, with a larger set of points than those provided to STalign, allowing an objective and fair comparison of geometric consistency across methods. This separation between optimisation and evaluation allows GALA to remain fully automated and reproducible, while still enabling an objective assessment of alignment quality.

### 3.5. Computational efficiency

All methods were benchmarked under consistent hardware conditions: if a baseline supports GPU computation, it was run on the same GPU as GALA; otherwise, it was executed on the same CPU. Details of the operating system and device configuration are provided in Supplementary Section S2.3. Across multiple benchmarks, GALA demonstrated superior computational efficiency, achieving both faster runtime and substantially lower memory usage compared with existing methods. It consistently outperformed other nonlinear methods, being substantially faster than GPSA and ST-GEARS (averaging 135.22 seconds compared with 245.61 and 154.69 seconds, respectively, on slices 2 and 3 of Sample C), while also maintaining a minimal memory footprint. While specialized rigid methods like PASTE, STAligner, and SLAT were faster (all under 30 seconds), they required substantially more memory and yielded inferior accuracy. Furthermore, GALA also outperformed STalign, its primary competitor built on the same LDDMM core, across multiple datasets. For example, on the human breast cancer dataset, GALA achieved an average runtime of 74.16 seconds with minimal memory usage, compared with STalign’s 512.06 seconds and substantially higher memory consumption, due to GALA’s streamlined optimisation and reduced parameters (see Section 2.3) .

GALA also excelled at scale where many methods failed due to excessive memory demands (e.g. 500 GB, MERFISH mouse liver data); it successfully aligned datasets of hundreds of thousands of cells in under a minute, far outpacing SLAT’s lengthy cell-matching phase. This efficiency extended to multimodal and mismatched-resolution scenarios, solidifying GALA’s advantage for rapid, low memory usage, high-fidelity alignment of large-scale ST data. The details of comparison of computational efficiency with baseline methods are provided at Supplementary Tab. S4.

## 4. Discussion

In this study, we introduced GALA, a unified, landmark-free spatial alignment framework capable of addressing a wide range of challenges in ST alignment. A central strength of GALA lies in its multimodal rasterisation strategy, which converts diverse transcriptomic and histological data into co-registered raster tensors. This standardised representation harmonises heterogeneous resolutions, modalities, and sequencing platforms, and naturally scales from spot-level to dense single-cell data, enabling the aggregation of millions of cells onto a user-defined grid where each pixel encodes local features. By design, GALA jointly leverages transcriptomic and histological information with tunable weights, enhancing both alignment accuracy and interpretability. Its coarse-to-fine optimisation couples a global affine transformation (GA), which identifies overlapping regions for initialisation, with an EM-like LDDMM refinement capable of large-scale deformations. The EM-style probabilistic matching focuses alignment on confidently corresponding regions while downweighting background or artefacts, making the coupled strategy robust across resolutions, modalities, and partial tissue overlaps. Furthermore, the invertibility of the diffeomorphic transformation ensures directionally consistent alignment. In within-resolution tasks, source-to-target and target-to-source alignments yield highly similar performance (Supplementary Section S2.2.1, Fig. S8). For cross-resolution or cross-modality alignment, the same property allows the high-resolution modality to be stably aligned to a lower-resolution reference, while the inverse transformation naturally provides the corresponding reverse mapping (Supplementary Fig. S16).

Compared with existing approaches (Tab. 1), GALA offers broader applicability and improved robustness. Spot-level methods such as PASTE and STAligner emphasise rigid alignment, whereas STalign and GPSA provide non-rigid deformations but with limited modality support or scalability. SLAT relies on one-to-one graph matching, which restricts its flexibility in heterogeneous tissue contexts. Coarse-to-fine frameworks such as ST-GEARS are specialised for particular modalities or task types, and most baselines, except for PASTE2, cannot handle partial tissue overlaps. In contrast, GALA uniquely integrates rigid and non-rigid deformations within a coupled framework and supports spot–to-spot, cell–to-cell, cell–to-spot, and transcriptomic–to-histology alignment across both full and partial tissue overlaps. While STalign also employs LDDMM, it cannot leverage gene expression or H&E image data, does not support spot-to-spot or direct cell-to-spot alignment, and relies on manually defined landmarks.

**Table 1.**
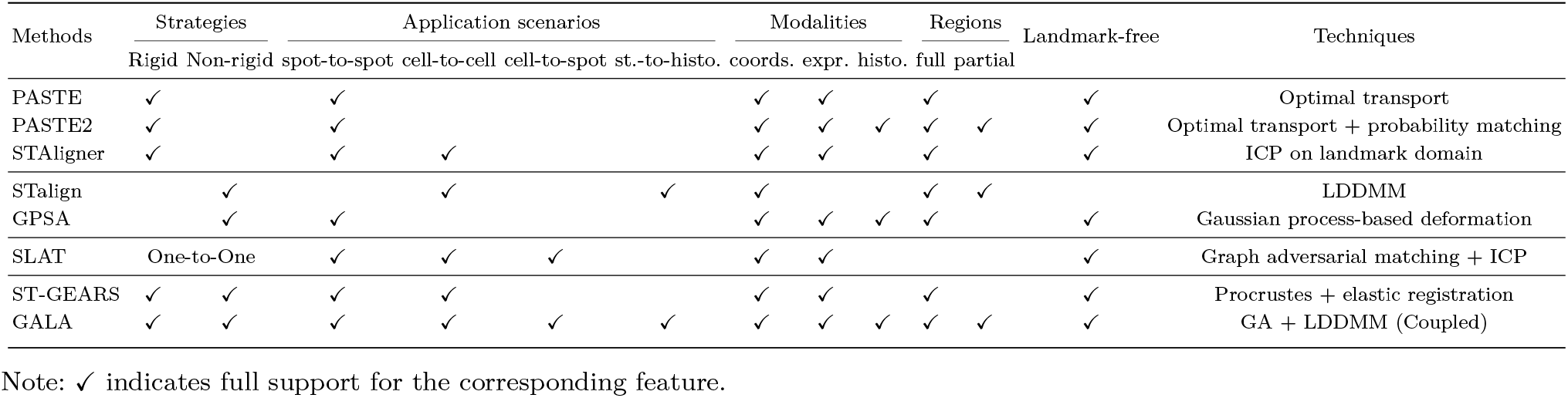
Summary of all baselines and GALA.

Despite its strengths, several limitations of GALA should be considered. Multimodal fusion of transcriptomic and histological data does not always improve alignment, reflecting both methodological constraints and the inherent biological reality that molecular and morphological states are not perfectly concordant [48]. Morphologically similar cells may exhibit distinct molecular profiles, and sharp molecular boundaries may lack histological counterparts[47]. Such discordances can introduce noise, highlighting the need for adaptive, biologically informed fusion strategies that weight each modality based on local context[3]. Furthermore, GALA focuses on geometric spatial alignment and does not explicitly correct gene expression batch effects. In datasets with substantial donor variability or platform differences, additional downstream integration methods may be required after alignment to harmonise molecular signals. Finally, as with all LDDMM-based frameworks, GALA assumes smooth, diffeomorphic deformations. Extremely irregular, discontinuous, or topology-disrupting distortions may violate this assumption and thus fall outside the model’s representational capacity, leading to alignment failure (Supplementary Section S2.2.2).

Nonetheless, by providing accurate and flexible spatial alignment, GALA establishes a critical foundation for downstream applications, including cross-sample atlas construction, consistent cell-type mapping, and trajectory inference. For example, GALA ensures spatial correspondence across samples, while network-based models (e.g. Perturb-STNet [17]) capture cell–cell interactions and perturbation effects within the aligned coordinate system, enabling integrative analyses that combine geometric alignment with functional modelling. Beyond transcriptomics, GALA is readily extendable to other spatially resolved omics modalities, such as spatial proteomics or metabolomics [49, 50]. By aggregating molecular measurements onto a common spatial grid through rasterisation, multi-omics data can be aligned within a unified framework. These representations may further be embedded into shared latent spaces using representation learning approaches, facilitating cross-modality comparison of spatial patterns while preserving modality-specific variation. Future work will explore joint multimodal alignment while addressing cross-modality consistency and heterogeneous noise. Taken together, integrating precise spatial alignment with downstream biological modelling represents a promising direction. We anticipate that GALA will serve as a versatile component within broader spatial omics pipelines, potentially coupled with modules for spatially informed gene expression modelling, or used to infer conserved spatial programmes across individuals, conditions, and technologies.

## 5. Data Availability

All datasets used in this study are publicly available from the original sources as cited below and were employed to benchmark GALA across various alignment tasks: The **DLPFC dataset**, used for full and partial spot-based alignment, was obtained from the Lieber Institute (https://github.com/LieberInstitute/HumanPilot/tree/master/10X); preprocessed data were downloaded from Zenodo (https://zenodo.org/records/6334774). The **mouse brain sagittal posterior dataset** (Sections 1 and 2), used to evaluate multimodal alignment and histology integration at the spot level, was downloaded from 10x Genomics: https://www.10xgenomics.com/datasets/mouse-brain-serial-section-1-sagittal-posterior-1-standard-1-1-0 and https://www.10xgenomics.com/datasets/mouse-brain-serial-section-2-sagittal-posterior-1-standard-1-1-0. The **MERFISH mouse liver dataset** (L2S1 and L2S2 from Animal 2), used for cell-level alignment without landmarks, was obtained from Vizgen: https://info.vizgen.com/mouse-liver-data. The **Visium mouse brain coronal section dataset**, used for cross-platform spot-to-cell alignment, was downloaded from: https://www.10xgenomics.com/datasets/adult-mouse-brain-coronal-section-fresh-frozen-1-standard. The **Xenium mouse brain dataset** (Replicate 2), used for cross-platform and cross-modality alignment with Visium: https://www.10xgenomics.com/datasets/fresh-frozen-mouse-brain-replicates-1-standard. The **Xenium human lung cancer FFPE dataset** (Experiment 1), used for evaluating image-to-cell alignment and multimodal registration in human tissue: https://www.10xgenomics.com/datasets/xenium-human-lung-cancer-post-xenium-technote. The **Xenium human breast cancer FFPE dataset** (Replicate 1), used for image-to-cell alignment and multimodal registration in human tissue: https://www.10xgenomics.com/products/xenium-in-situ/preview-dataset-human-breast.

All preprocessed ST datasets and corresponding alignment outputs generated in this study are publicly available via Zenodo (https://zenodo.org/records/17576488) to allow full reproduction of the analyses and figures presented.

## 6. Code Availability

The source code implementing `GALA` is openly available at https://github.com/TaoDing2/GALA under an open-source licence. The repository provides detailed instructions for installation, data preparation, and reproduction of all analyses. In addition, the Zenodo repository contains scripts and processed data necessary to replicate all experiments and figures reported in this study.

## 8. Author contributions

T.D. and P.Z. conceived the study. T.D. developed the GALA algorithm, performed data analyses, and drafted the manuscript. P.Z. supervised the project, provided conceptual guidance, and contributed to manuscript revisions. All authors read and approved the final manuscript.

## 9. Biographical Note

T.D. and P.Z. are researchers in computational biology and spatial omics, focusing on multimodal data integration and the development of bioinformatics methods.

