## Supplement for "GALA: A Unified Landmark-Free Framework for Coarse-to-Fine Spatial Alignment Across Resolutions and Modalities in Spatial Transcriptomics"

### S1 Methodological details

#### S1.1 Rasterisation examples at varying grid spacings

Each dataset is represented as a co-registered tensor, with the source $I \in \mathbb{R}^{c \times H_1 \times W_1}$ and the target $J \in \mathbb{R}^{c \times H_2 \times W_2}$. For transcriptomic-only inputs ($c = p+1$), $I = I_{\text{st}}$ and $J = J_{\text{st}}$, and when histology is included ($c = p+2$), $I = (I_{\text{st}}, I_{\text{histo}})$ and $J = (J_{\text{st}}, J_{\text{histo}})$, with concatenation along the channel dimension. The spatial dimensions $(H_1, W_1)$ and $(H_2, W_2)$ may differ according to each dataset's native resolution $dx$ and field of view. This unified raster representation provides a common interface for all subsequent optimisation.

To illustrate the effect of grid resolution $dx$, we visualised rasterised expression and histological features at two spatial scales using the mouse brain sagittal–posterior (MBSP) Visium sections and the MERFISH mouse liver dataset. In the MBSP samples (platform-provided *lowres* images), $dx \approx 1$ unit preserved sharp spot-level variation, whereas $dx \approx 5$ units generated smoother, more diffuse patterns (Fig.S1a). In the MERFISH liver data, $dx = 30$ $\mu m$ retained fine-grained spatial patterns, while $dx = 100$ $\mu m$ produced a coarser aggregated representation (Fig.S1b). Across both settings, smaller $dx$ values preserve fine-scale structure, whereas larger $dx$ emphasise broader spatial trends.

Throughout this study, we use $dx \approx 1$ unit for spot-level data and $dx = 30$ $\mu m$ for cell-level data. This choice balances spatial fidelity and computational efficiency, and can be adapted according to the requirements of downstream alignment or analysis tasks. Details of the sensitivity analysis of grid spacing $dx$ are provided in Section S2.1.3.

#### S1.2 Objective function

**Coarse alignment.** We initialise the alignment by estimating a global affine transformation between the source and target domains. This transformation is parametrised by five variables: rotation $\theta$, translation $(t_x, t_y)$, and independent scaling factors $(s_x, s_y)$. Reflections are naturally accommodated by allowing negative scale parameters, which frequently arise in cross-modality settings due to mirrored or flipped tissue orientations. The affine transformation is defined as

$$A(\boldsymbol{x}) = \left( \underbrace{\begin{bmatrix} \cos\theta & -\sin\theta & 0 \\ \sin\theta & \cos\theta & 0 \\ 0 & 0 & 0 \end{bmatrix}}_{\text{Rotation}} \underbrace{\begin{bmatrix} \pm s_x & 0 & 0 \\ 0 & \pm s_y & 0 \\ 0 & 0 & 0 \end{bmatrix}}_{\text{Scaling \& Reflection}} + \underbrace{\begin{bmatrix} 0 & 0 & t_x \\ 0 & 0 & t_y \\ 0 & 0 & 1 \end{bmatrix}}_{\text{Shift}} \right) \begin{bmatrix} x \\ y \\ 1 \end{bmatrix},$$

where $\boldsymbol{x} = (x, y, 1)^T$ denotes the 2D raster coordinate in homogeneous form, and $A(\boldsymbol{x})$ gives its affine-transformed position.

**Fine alignment.** Following coarse alignment, we refine the correspondence using large deformation diffeomorphic metric mapping (LDDMM)(Beg et al., 2005), which models non-linear deformations via a smooth, invertible deformation $\phi^v$ that preserves tissue topology. This formulation is well suited to biological samples, where local distortions arise from deformation, processing, or sectioning.

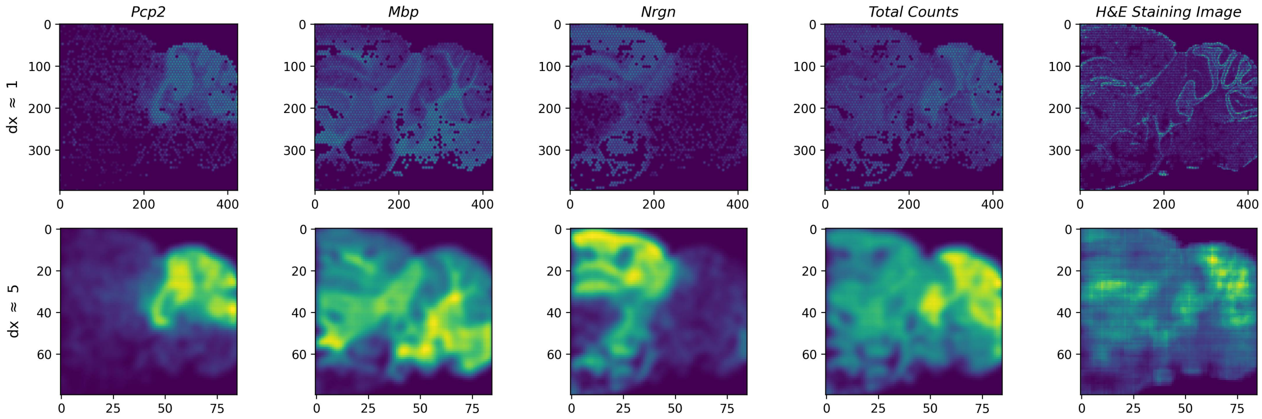

(a) MBSP sections

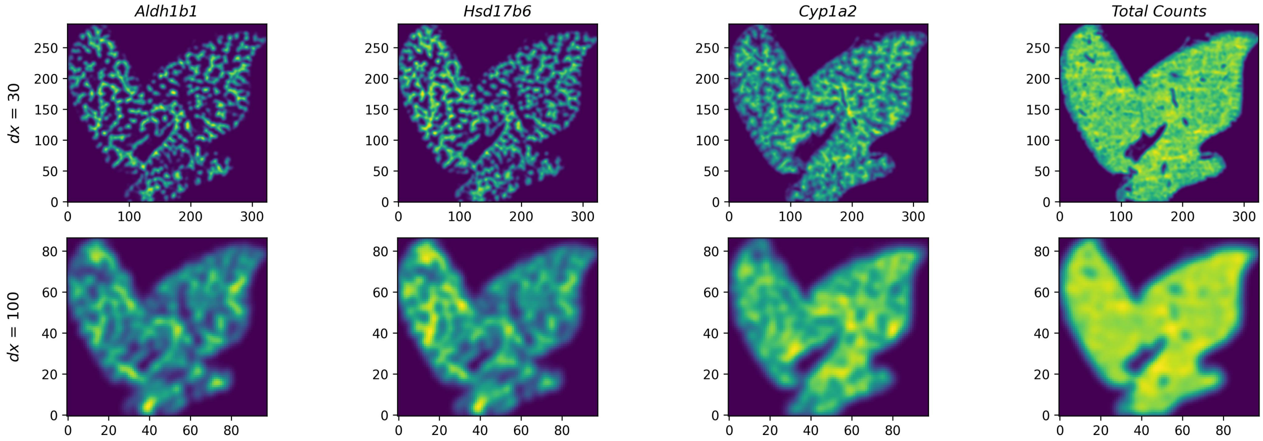

(b) Mouse liver map

Figure S1: **Examples of rasterised transcriptomic and histological features (when available) under varying grid spacings**. Each column corresponds to one channel, including three selected genes, total expression counts, and the H&E image. Brighter colours indicate higher values after rasterisation. All expression channels are normalised to $[0, 1]$ for visual comparability. Increasing $dx$ progressively smooths spatial structure, producing lower-resolution yet more continuous representations. **(a)** Visium-based MBSP dataset at spot-level resolution. The top row shows rasterised tensors at a fine scale ($dx \approx 1$ unit), while the bottom row shows coarser outputs at $dx \approx 5$ units. **(b)** MERFISH-based mouse liver dataset at cell-level resolution. The two rows correspond to raster grids of $dx = 30\ \mu m$ and $dx = 100\ \mu m$, respectively.

Let $v_t(\boldsymbol{x})$ denote a time-dependent velocity field for $t \in [0, 1]$. Integration of $v_t$ yields a diffeomorphic flow $\phi_t^v(\boldsymbol{x})$ satisfying the ordinary differential equation:

$$\frac{d\phi_t^v(\boldsymbol{x})}{dt} = v_t(\phi_t^v(\boldsymbol{x})), \quad \phi_0^v(\boldsymbol{x}) = \boldsymbol{x}. \tag{S1}$$

The subscript explicitly denotes the dependence of $\phi$ on the associated velocity field $v$. The initial velocity field is determined from the coarse affine configuration, ensuring a coupled progression from global initialisation to local refinement. At $t = 0$, $\phi_0^v = Id$ is the identity map, while $\phi_1^v \doteq \phi^v$ at $t = 1$ denotes the final diffeomorphism linking the datasets such that $J(\boldsymbol{x}) \approx I(\phi^v(\boldsymbol{x}))$.

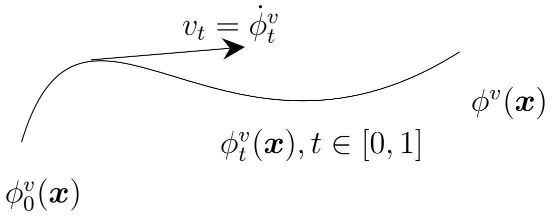

Figure S2: **Velocity field underlying diffeomorphic mapping**.

**Probabilistic matching framework.** Although the affine component captures coarse geometry and accommodates partial tissue overlap, and the diffeomorphic component accounts for smooth local deformation, neither alone can fully address local disruptions such as tears, folds, or modality-specific artefacts. To model these irregularities, we introduce a probabilistic formulation that distinguishes between regions with reliable biological correspondence (*matched*) and those without (*unmatched*). Prior EM-like LDDMM approaches (Tward et al., 2020; Clifton et al., 2023) have used three or more latent classes; however, we adopt a two-class model for three reasons: (i) background and artefactual regions generally lack meaningful correspondence; (ii) upstream gene selection and rasterisation already downweight low-quality areas; and (iii) in multimodal alignment, structural consistency is more relevant than explicit pixel-level pairing.

Let $I(\boldsymbol{x}), J(\boldsymbol{x}) \in \mathbb{R}^c$ denote the raster intensities of source and target at location $\boldsymbol{x} \in \Omega$. We assume that target intensities at location $\boldsymbol{x}$ is

$$J(\boldsymbol{x}) = \begin{cases} I((\phi^v \cdot A)(\boldsymbol{x})) + \boldsymbol{\epsilon}_M, & \boldsymbol{x} \text{ is matched,} \\ \boldsymbol{\mu}_N + \boldsymbol{\epsilon}_N, & \boldsymbol{x} \text{ is unmatched,} \end{cases}$$

where the residuals $\boldsymbol{\epsilon}$ are defined as deviations from their respective mean intensities. To quantify similarity, we evaluate these residuals using Gaussian kernels

$$K(J, \boldsymbol{\mu}, \Sigma) = \exp\Big( - \frac{1}{2}(J - \boldsymbol{\mu})^\top \Sigma^{-1} (J - \boldsymbol{\mu})\Big)$$

with covariances $\Sigma_M = \sigma_M^2 I_c$ and $\Sigma_N = \sigma_N^2 I_c$, and $I_c$ is a $c \times c$ identity matrix. $\boldsymbol{\mu}_N$ denotes the mean intensity of non-matching regions, estimated iteratively. This Gaussian form induces a quadratic data term and enables closed-form EM updates; it is used as a variational similarity construction rather than a strict distributional assumption. Given class priors $\pi_M$ and $\pi_N$ ($\pi_M + \pi_N = 1$), the posterior probability that location $\boldsymbol{x}$ belongs to the matched class is computed as

$$P_M(\boldsymbol{x}) = \frac{\pi_M \, K(J(\boldsymbol{x}), I((\phi^v \cdot A)(\boldsymbol{x})), \Sigma_M)}{\pi_M \, K(J(\boldsymbol{x}), I((\phi^v \cdot A)(\boldsymbol{x})), \Sigma_M) + \pi_N \, K(J(\boldsymbol{x}), \boldsymbol{\mu}_N, \Sigma_N)}. \tag{S2}$$

The posterior map $P_M(\boldsymbol{x})$ acts as a soft indicator of reliable alignment and is updated jointly with the deformation parameters.

**Objective function** Accordingly, we define the objective function as shown in Eq. (1) of the main manuscript, reproduced here for completeness:

$$\begin{aligned}\mathcal{L}(A, \phi^v, P_M) = {} & \alpha \frac{1}{2\sigma_{M_1}^2} \sum_{g=1}^{p+1} \sum_{\boldsymbol{x}} P_M(\boldsymbol{x}) \, \|I_{\text{st}}^g((\phi^v \cdot A)(\boldsymbol{x})) - J_{\text{st}}^g(\boldsymbol{x})\|_2^2 \\ & + (1-\alpha) \frac{1}{2\sigma_{M_2}^2} \sum_{\boldsymbol{x}} P_M(\boldsymbol{x}) \, \|I_{\text{histo}}((\phi^v \cdot A)(\boldsymbol{x})) - J_{\text{histo}}(\boldsymbol{x})\|_2^2 \\ & + \frac{1}{2\sigma_R^2} \int_0^1 \|v_t\|_V^2 dt \end{aligned} \tag{S3}$$

The detailed parameter demonstration are listed below:

(1). $I_{\text{st}}^g$ and $J_{\text{st}}^g$ denote the $g$-th channel of the source and target transcriptomic raster tensors, where $g = 1, \ldots, p$ correspond to selected informative genes and $g = p + 1$ represents the total gene expression;

(2). $I_{\text{histo}}$ and $J_{\text{histo}}$ are the source and target histological raster tensor, co-aligned with transcriptomic tensor.

(3). $\alpha \in (0, 1)$ is the fusion weight balancing transcriptomic and histological features.

(4). $\|\cdot\|_2$ represents $L^2$ norm.

(5). The velocity vector field is regularized by a norm $\|\cdot\|_V$ in the space of vectors fields $V$ (Beg et al., 2005) with the inner product defined through a differential operator $L$ (denoting its adjoint as $L^\dagger$) given by

$$\langle f, g \rangle_V \doteq \langle L^\dagger L f, g \rangle_{L^2}.$$

In applications of Tward et al. (2020) and Clifton et al. (2023), the operator $L$ is chosen to be of the Cauchy-Navier type, and is self-adjoint (i.e., $L = L^\dagger$), as $L = (I - a^2 \Delta)^2$, where

(a). $a$ is a constant with the units of length that controls spatial smoothness.

(b). $\Delta$ is the Laplacian operator.

(c). The power of $L$ (set as 2) is to guarantee that results are diffeomorphisms (Dupuis et al., 1998).

Hence, the regularisation term in the cost function can be represented as

$$\begin{aligned}\int_0^1 \|v_t\|_V^2 dt &\doteq \int_0^1 LLv_t \cdot v_t dt = \int_0^1 \|Lv_t\|_2^2 dt \\ &= \int_0^1 \int_{\mathbb{R}^2} \|(I - a^2\Delta)^2 v_t\|_2^2 dt\end{aligned}$$

(6). $\sigma_{M_1}, \sigma_{M_2}, \sigma_R, \sigma_N$ are user tunable parameters, controlling the relative influence of each component and scaling of each term.

Detailed descriptions of the default parameters are provided in Tab. S1.

#### S1.3 Optimisation and convergence of GALA

In the objective function defined in Eqn. S3, it is natural to consider jointly estimating the affine transformation, the non-linear deformation, and the class assignment of each spots/cells. Indeed, previous studies (Tward et al., 2020; Clifton et al., 2023) have proposed incorporating affine transformations within the LDDMM framework via a non-monotonic polynomial mapping $f_\theta$, for example $f_\theta(I(\phi(\boldsymbol{x}))) = a(I(\phi(\boldsymbol{x}))) + b$ when $\theta = 1$. However, in these approaches the parameters are updated by minimising a data-fidelity term of the form $\frac{1}{2\sigma^2}|aI(\phi(\boldsymbol{x})) + b - J(\boldsymbol{x})|_2^2$ at each iteration, which makes the results highly sensitive to the initial estimates of $a$ and $b$, thereby compromising robustness.

In practice, jointly optimising both global affine transformation and local diffeomorphic deformation can be challenging. These two components serve distinct geometric functions: affine transformations account for coarse, global misalignments such as rotations, translations, scaling, and reflections, whereas LDDMM captures smooth, local, non-linear deformations. Simultaneous estimation can lead to competition between the two components in explaining overlapping spatial discrepancies, resulting in unstable convergence and potential overfitting. Additionally, the enlarged parameter space increases computational cost and renders the optimisation landscape more complex, complicating gradient-based updates. Finally, interpretability is reduced, as it becomes unclear whether a given spatial correspondence arises from global warping or local deformation.

To address these challenges, GALA employs a coupled coarse-to-fine optimisation strategy. Specifically, global alignment is first achieved using a GA-based population search over the affine parameters (Gad, 2024), which effectively handles large initial displacements and efficiently identifies the overlap region between source and target. Building on this configuration, LDDMM subsequently estimates a smooth, invertible deformation through a time-dependent velocity field. The refinement follows an EM-like procedure (Tward et al., 2020; Clifton et al., 2023): match probabilities are updated in the E-step, while the velocity field is updated in the M-step by minimising the weighted objective function.

Unlike conventional coarse-to-fine pipelines where global and local stages are loosely coupled, GALA integrates the two components under a single objective function (Eqn. S3). This coupling ensures that (i) large-scale misalignments are robustly corrected before local refinement, (ii) uncertainty estimates propagate consistently from coarse to fine scales, and (iii) alignment focuses on reliable regions while automatically downweighting uncertain or unmatched areas. Collectively, these advantages improve both the robustness and interpretability of the alignment results. The complete optimisation procedure is summarised in pseudocode in Alg. S1, with default parameters listed in Tab. S1.

Table S1: Default parameters used in GALA

| Parameter | Description | Default |
|---|---|---|
| $\alpha$ | Weighting between gene expression and histology features | 0.5 |
| $p$ | Number of top predictable genes | 3 |
| *num_generations* | Number of generations | 1000 |
| *num_parents_mating* | Number of solutions selected as parents | 50 |
| *sol_per_pop* | Population size (number of solutions) | 100 |
| *mutation_probability* | Probability of mutation | 0.2 |
| *crossover_probability* | Probability of selecting a parent for crossover | 0.8 |
| *random_seed* | Random seed | None |
| $\theta$ | Rotation angle (positive: counter-clockwise; negative: clockwise) | $[-\pi, \pi]$ |
| $t_x$ | Translation along $x$-axis | $\left[-\frac{r_{Jx}}{2},\ \frac{r_{Jx}}{2}\right]$ |
| $t_y$ | Translation along $y$-axis | $\left[-\frac{r_{Jy}}{2},\ \frac{r_{Jy}}{2}\right]$ |
| $s_x$ | Scaling along $x$-axis (negative: horizontal reflection) | $[-(s_x^{\text{range}}+0.1),\ (s_x^{\text{range}}+0.1)]$ |
| $s_y$ | Scaling along $y$-axis (negative: vertical reflection) | $[-(s_y^{\text{range}}+0.1),\ (s_y^{\text{range}}+0.1)]$ |
| *num_iterations* | Number of iterations | 5000 |
| $a$ | Smoothness scale of diffeomorphism | $dx$ (raster grid spacing) |
| $\Delta_{v_t}$ | Gradient descent step size for $v_t$ | $dx$ |
| $\sigma_{M_1}, \sigma_{M_2}$ | Weight on matching region | Standard deviation of $I$; $\sigma_{M_1} = \sigma_{M_2}$ |
| $\sigma_B$ | Weight on background term | Standard deviation of $J$ |
| $\sigma_R$ | Weight on regularisation term | $1e3$ |

**Note:** $r_{Ix} = \max(x_I^{(x)}) - \min(x_I^{(x)})$, $r_{Iy} = \max(x_I^{(y)}) - \min(x_I^{(y)})$, $r_{Jx} = \max(x_J^{(x)}) - \min(x_J^{(x)})$, $r_{Jy} = \max(x_J^{(y)}) - \min(x_J^{(y)})$; $s_x^{\text{range}} = \max\left(\frac{r_{Ix}}{r_{Jx}}, \frac{r_{Jx}}{r_{Ix}}\right)$, $s_y^{\text{range}} = \max\left(\frac{r_{Iy}}{r_{Jy}}, \frac{r_{Jy}}{r_{Iy}}\right)$.

**Algorithm S1** Optimisation framework of GALA

**Input Data:** Source rasterised tensor $I \in \mathbb{R}^{c \times H_1 \times W_1}$ and target rasterised tensor $J \in \mathbb{R}^{c \times H_2 \times W_2}$.

**Input Parameters:** Regularisation parameters $\alpha, \sigma_{M_1}, \sigma_{M_2}, \sigma_N, \sigma_R$, and kernel width $a$.

**Initialization:**

1: Initialise affine matrix $A = I_3$.

2: Initialise diffeomorphism $\phi^v = Id$ by setting the velocity field $v_t = 0$ with 3 time steps $t \in [0, 1]$.

3: Initialise priors $\pi_M = \pi_N = 0.5$ and posterior maps $P_M(\boldsymbol{x}), P_N(\boldsymbol{x})$.

4: Precompute the differential operator

$$LL = \left(1 + 2a^2 \sum_{i=1}^{2} \frac{1 - \cos(2\pi \Delta_{x_i} k_i)}{\Delta_{x_i}^2}\right)^{2*2},$$

where $\Delta_{x_i}$ is the grid step along axis $i$ and $k_i$ denotes the corresponding frequency component (Beg et al., 2005).

**Optimisation:** Repeat until the maximum number of outer episodes is reached.

**Affine update via GA (coarse alignment)**

1: Fix $\phi^v$.

2: Repeat until the GA stopping criterion is met:

(a) Maximise the objective $\frac{1}{\mathcal{L}(A, \phi^v, P_M)}$ via global search.

(b) Update affine parameters $(\theta, t_x, t_y, s_x, s_y)$ and form the matrix $A$.

**EM-like LDDMM update (fine alignment)**

3: Fix $A$.

4: Repeat until the LDDMM stopping criterion is met:

i. **E-step:** Update posteriors $P_M(\boldsymbol{x})$ and $P_N(\boldsymbol{x})$ via Eq. (S2), and set

$$\pi_M = \sum_{\boldsymbol{x}} P_M(\boldsymbol{x}), \pi_N = \sum_{\boldsymbol{x}} P_N(\boldsymbol{x})$$

ii. **M-step:**

(a) **Forward pass:** Integrate the velocity field using Eq. (S1) to obtain $(\phi_t^v \cdot A)(\boldsymbol{x})$; update the non-matching mean

$$\boldsymbol{\mu}_N = \frac{\sum_{\boldsymbol{x}} J(\boldsymbol{x}) P_N(\boldsymbol{x})}{\sum_{\boldsymbol{x}} P_N(\boldsymbol{x})}.$$

(b) **Objective evaluation:** Compute $I((\phi^v \cdot A)(\boldsymbol{x}))$ and evaluate $\mathcal{L}(A, \phi^v, P_M)$.

(c) **Backward pass:** Update the velocity field $v_t$ via gradient descent with smoothing by $LL^{-1}$.

**Output:** Affine parameters $(\theta, t_x, t_y, s_x, s_y)$, diffeomorphic deformation $\phi^v$, and posterior matching probabilities $P_M$.

We further examined the convergence behaviour of GALA during the first four episodes (each episode includes 1000 generations in GA and 5000 iterations in LDDMM) by aligning slice 3 to slice 2 of Sample C in the DLPFC dataset under the settings ($p = 3, \alpha = 0.5$), with a random seed of 42 and all other parameters as specified in Tab. S1. Notably, the optimal affine transformation was identified within this initial run and remained unchanged in subsequent episodes, during which only the LDDMM refinement continued to update the alignment. Fig. S3b shows the objective function across all four episodes, with the $x$-axis representing the cumulative number of iterations. Each episode builds on the previous one, yielding a total of 20,000 iterations.

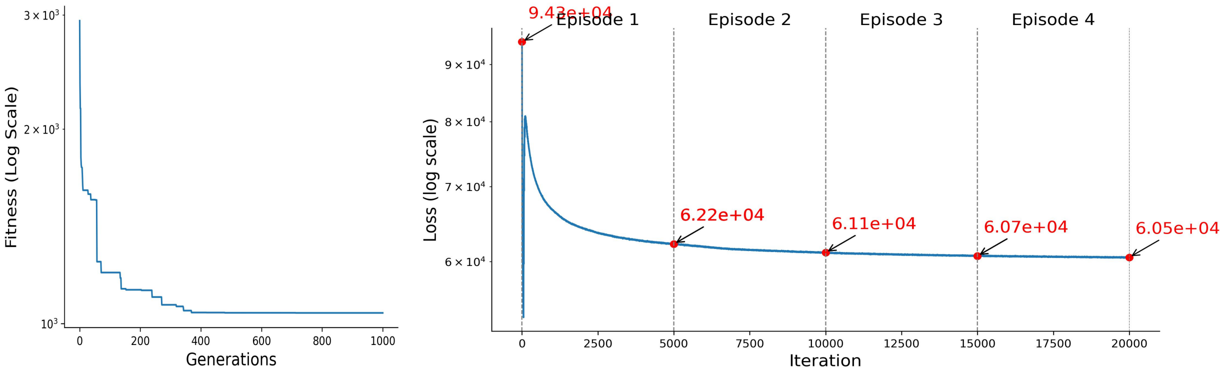

(a) Fitness over generations

(b) Objective function across four episodes

Figure S3: **Convergence behaviour of GALA on slices 2 and 3 of Sample C in the DLPFC dataset**. Both $y$-axes are log-scaled. **(a)** Fitness values during the first genetic algorithm episode, indicating that the optimal affine parameters are typically found early. **(b)** Loss values over 20,000 iterations, segmented into four consecutive episodes as annotated at the top of the figure. Red dots mark the objective values at the beginning of the first episode and at the end of each episode.

The gradient descent step size $\Delta_{v_t}$ exerts a significant influence on convergence speed. In our experiments, $\Delta_{v_t}$ was generally set to match the unit grid spacing, $dx$, ensuring stable optimisation. As shown in Fig. S3, the objective function decreases sharply during the first two episodes but plateaus thereafter, with further iterations incurring substantial computational cost while providing only marginal gains in alignment accuracy. Accordingly, we limited our analyses to two episodes across all experiments in this study.

#### S1.4 Evaluation metrics

**Label-based evaluation.** We introduce a coordinate-based label consistency accuracy to quantitatively assess the effectiveness of GALA alignment. Following alignment, mutual nearest neighbour (MNN) spots/cells pairs are identified using Euclidean distances between spatial coordinates in the aligned space. For each spot in the source slice, its nearest neighbour in the target slice is located, and vice versa; only reciprocal matches are retained. Alignment accuracy is then defined as the proportion of MNN pairs whose constituent spots share the same annotated cell type. Formally:

$$\text{Accuracy} = \frac{1}{N}\sum_{i=1}^{N} \mathbb{1}\left[\text{type}(\boldsymbol{s}_i) = \text{type}(\boldsymbol{t}_i)\right] \tag{S4}$$

where $(\boldsymbol{s}_i, \boldsymbol{t}_i)$ denotes the $i$th MNN pair, with $\boldsymbol{s}_i$ and $\boldsymbol{t}_i$ representing aligned coordinates in the source and target slices, and $N$ the total number of identified pairs. This metric captures the degree to which biologically similar spots are brought into correspondence through spatial alignment.

We additionally evaluate alignment by fitting a Gaussian mixture model (GMM) to the concatenated source and target data, incorporating both gene expression and spatial information. Clustering quality is then assessed using standard metrics such as the Adjusted Rand Index (ARI) and Normalised Mutual Information (NMI), which quantify agreement with known cell type annotations. Higher ARI or NMI indicates that the aligned slices form coherent, biologically meaningful clusters.

**Spot-based evaluation.** For datasets without reliable anatomical labels, we quantify alignment fidelity using a spatial cross-correlation (SCC) score computed for each gene. Let $\boldsymbol{y}^{(1)}, \boldsymbol{y}^{(2)} \in \mathbb{R}^{n\times p}$ denote the expression matrices for two slices, with $n$ spots and $p$ genes, and let $\boldsymbol{x}_i^{(1)}, \boldsymbol{x}_j^{(2)} \in \mathbb{R}^2$ denote the corresponding spatial coordinates.

We first compute the Euclidean distance matrix:

$$d_{ij} = \|\boldsymbol{x}_i^{(1)} - \boldsymbol{x}_j^{(2)}\|_2$$

and convert it into a spatial weight matrix $W_{ij} = \exp(-\beta d_{ij}^2)$, where $\beta > 0$ controls the spatial decay. Let $S_{ij} = \sum_{i=1}^{n}\sum_{j=1}^{n} W_{ij}$.

For gene $g$, with expressions $\boldsymbol{y}_i$ and $\boldsymbol{z}_j$ in slices 1 and 2 and means $\bar{\boldsymbol{y}}$ and $\bar{\boldsymbol{z}}$, the SCC score is defined as

$$\text{SCC}_g = \frac{n}{S_{ij}} \cdot \frac{\sum_{i=1}^{n}\sum_{j=1}^{n} W_{ij}\,(\boldsymbol{y}_i - \bar{\boldsymbol{y}})(\boldsymbol{z}_j - \bar{\boldsymbol{z}})}{\sqrt{\sum_{i=1}^{n}(\boldsymbol{y}_i - \bar{\boldsymbol{y}})^2}\,\sqrt{\sum_{j=1}^{n}(\boldsymbol{z}_j - \bar{\boldsymbol{z}})^2}}.$$

This extends the Pearson correlation by weighting cross-slice spot pairs according to spatial proximity. A high $\text{SCC}_g$ indicates strong preservation of the spatial expression pattern of gene $g$ across slices following alignment. Scores may be averaged across genes or examined individually to identify the most spatially conserved molecular features.

### S2 Supplementary analyses of GALA

#### S2.1 Robustness and sensitivity analyses of key hyper-parameters

The objective function (Eqn. S3) comprises three components: a transcriptomic fidelity term, a histological fidelity term, and a regularisation term. The transcriptomic fidelity term combines total gene expression counts with the top $p$ spatially informative genes, identified via $R^2$ (Sec. 4.3). The histological fidelity term captures structural cues from tissue morphology, such as edges and local textures, providing complementary information to transcriptomic measurements. The two fidelity terms are jointly weighted by the fusion parameter $\alpha$, which determines the relative contribution of molecular and morphological features. The regularisation term follows established LDDMM formulations (Tward et al., 2020; Clifton et al., 2023), encouraging smooth deformations while preventing overfitting.

To evaluate the robustness and sensitivity of parameters in GALA and fidelity terms, we first examined the effect of the gene number $p$ under varying degrees of overlap in partial alignment tasks (Section S2.1.1). We then quantified the joint influence of the fusion weight $\alpha$ and the number of selected genes $p$ (Section S2.1.2), thereby characterising the stability of GALA across a broad parameter range. Third, we examined the sensitivity of alignment performance to the choice of $dx$, which determines the spatial resolution of the rasterisation (Section S2.1.3). Lastly, we considered cases where the selected informative genes exhibit substantial spatial overlap or incomplete tissue coverage (Section S2.1.4).

Unless otherwise specified, all experiments used a random seed of 42 for reproducibility. The GA was run for 1,000 generations, followed by LDDMM with a maximum of 5,000 iterations. Owing to early convergence of the objective function (Fig. S3b), only two optimisation episodes were required.

##### S2.1.1 Robustness to the number of informative genes under varying overlap conditions

We quantified how the number of spatially informative genes influences alignment performance under varying overlap ratios using transcriptomics-only inputs. Specifically, we examined $p \in \{0, 1, 2, 3, 4, 5, 7, 10, 15\}$, where $p$ denotes the number of selected informative genes ranked by $R^2$. The case $p = 0$ uses only total gene expression counts. For each overlap ratio $s \in \{0.3, 0.5, 0.6, 0.7, 0.8, 0.9, 1.0\}$, we aligned a left subregion of slice 3 from Sample C to the full slice 2, with the retained area representing the overlap. Label consistency accuracy was computed for each configuration.

As shown in Fig. S4, values of $p$ = 1–3 offered the best balance between stability and accuracy across conditions. The control case ($p = 0$) performed poorly, even at 80% overlap. Larger gene sets ($p \geq 4$) frequently introduced redundancy or noise, reducing accuracy. Among these, $p = 3$ consistently achieved strong performance (0.83–0.88), making it the most robust choice across scenarios.

Together, these results show that a small set of carefully selected spatially informative genes is sufficient for robust alignment, even with limited overlap. Accordingly, we fixed $p = 3$ for all subsequent analyses, balancing accuracy and computational efficiency.

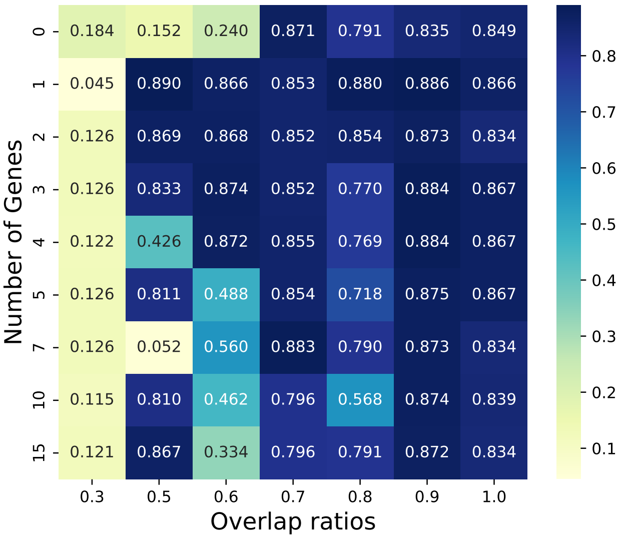

Figure S4: **Impact of the number of informative genes on spatial alignment accuracy across overlap ratios**. Performance is shown for varying numbers of spatially predictable genes ($p$, rows) across overlap ratios (columns, 30–100%). The control condition ($p = 0$, top row) uses only total expression counts without gene-specific channels.

##### S2.1.2 Sensitivity of performance to the fusion weight $\alpha$ and gene number $p$

We next conducted controlled perturbation studies on three human DLPFC samples (A, B, and C), aligning slice 3 to slice 2 in each case, and on the MBSP dataset. To assess sensitivity to the fusion weight, we fixed $p = 3$ and varied $\alpha \in \{0.1, 0.3, 0.5, 0.7, 0.9, 1\}$. To analyse sensitivity to the number of informative genes, we fixed $\alpha = 0.5$ and varied $p$ over the same set used above. These experiments characterise how alignment behaviour varies when deviating from nominal hyper-parameter settings.

For the DLPFC samples (Fig. S5a), alignment accuracy generally improves from $\alpha = 0.1$ to 0.7, with Samples B and C plateauing or peaking between $\alpha = 0.7$ and 1.0. Sample A displays a more variable trend, achieving its highest accuracy at $\alpha = 1.0$. These results indicate that moderate incorporation of histology is beneficial, whereas very small $\alpha$ values underutilise informative structural cues. In the $p$-sensitivity analysis ($\alpha = 0.5$ fixed), Samples B and C perform well even at $p = 0$, with marginal gains beyond $p = 3$. In Sample A, accuracy peaks at $p =$ 1–2 and decreases slightly as more genes are added. Solid curves show post-alignment performance, whereas dashed lines indicate the no-alignment baselines.

In the MBSP dataset (Fig. S5b), SCC scores increase as $\alpha$ increases towards 0.5, suggesting that moderate integration of histology improves alignment. For $\alpha \geq 0.5$, SCC values stabilise, indicating that transcriptomics alone provides strong spatial signals, with histology contributing marginal additional benefit. Under $\alpha = 0.5$, SCC remains stable across $p$, with slightly higher variability for $p > 5$, likely reflecting the noise introduced by including less informative genes. Overall, GALA shows strong robustness to both parameters, maintaining reliable performance even with very limited molecular input.

In summary, although performance remains stable across wide parameter ranges, we adopt $\alpha = 0.5$ and $p = 3$ as default choices. This configuration balances transcriptomics and histological contributions, aligns with the principle that both modalities should inform the

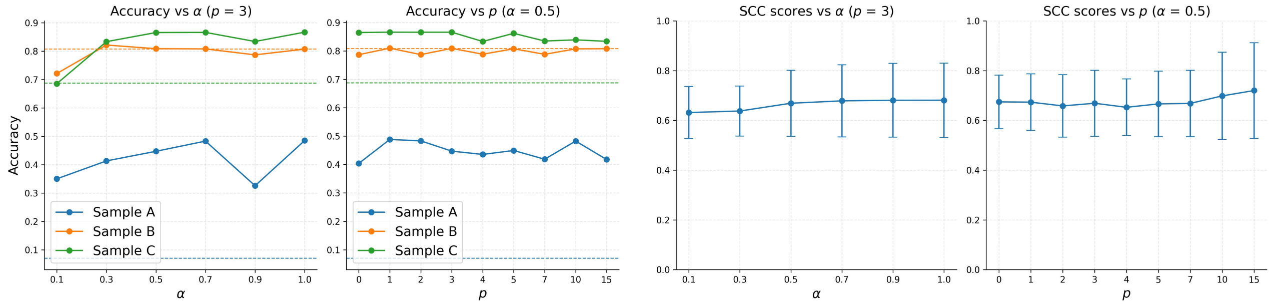

(a) DLPFC dataset (b) MBSP dataset

Figure S5: **Parameter sensitivity analysis with respect to fusion weight $\alpha$ and the number of informative genes $p$.** **(a)** Label consistency accuracy on three DLPFC samples (A, B, and C), aligning slice 3 to slice 2 in each case. Left: Accuracy across values of $\alpha$ (with $p = 3$ fixed). Right: Accuracy across values of $p$ (with $\alpha = 0.5$ fixed). Solid lines show post-alignment accuracy; dashed lines indicate baselines. **(b)** SCC scores on the MBSP sections. Left: SCC across $\alpha$ (with $p = 3$ fixed). Right: SCC across $p$ (with $\alpha = 0.5$ fixed). Each value is averaged over 135 genes with standard errors.

objective comparably, and generalises well across datasets.

##### S2.1.3 Sensitivity analysis of grid spacing $dx$

Section S1.1 illustrated two rasterisation examples using different grid spacings $dx$ for MBSP sections and mouse liver replicates. Here, we further examine the sensitivity of alignment performance to the choice of $dx$. The grid spacing $dx$ determines the spatial resolution of the rasterised representation used in intensity-based matching. Smaller $dx$ yields finer spatial discretisation, potentially capturing more local structural detail but increasing computational cost. Larger $dx$ reduces resolution and computational burden, at the risk of losing fine-scale spatial variation.

Because the DLPFC dataset provides reliable anatomical annotations, it enables quantitative evaluation of alignment accuracy in addition to computational cost. Therefore, we performed sensitivity analysis under varying grid spacings (Tab. S2) on two representative datasets: Sample C (slice 2 and slice 3, spot-level) of the DLPFC dataset and the mouse liver map dataset (cell-level). For the DLPFC data, we recorded label consistency accuracy, whereas for the mouse liver data, we measured cosine similarity between pseudo-spots. For both datasets, runtime and GPU memory consumption were also recorded. All experiments were conducted with GPU acceleration.

Across both datasets, alignment accuracy exhibits a mild U-shaped dependence on $dx$, consistent with the expected trade-off between spatial resolution and discretisation sparsity. When $dx$ is excessively small, the rasterised representation becomes increasingly sparse and computationally demanding, without yielding substantial improvement in alignment accuracy. Conversely, when $dx$ is overly large, spatial resolution is reduced and fine-scale structural information is gradually lost, leading to modest performance degradation.

Importantly, within a broad intermediate range around the selected default values, alignment accuracy varies only mildly. This relative robustness arises from the continuous deformation formulation of LDDMM, where smoothness regularisation on the velocity field stabilises

Table S2: Sensitivity analysis with respect to grid spacing $dx$

| $dx$ | 0.450 | 0.675 | 0.900 | 1.125 | 1.350 | 2.250 | 4.500 |
|---|---|---|---|---|---|---|---|
| Accuracy | 0.833 | 0.864 | 0.867 | 0.846 | 0.844 | 0.830 | 0.819 |
| Runtime (s) | 625.657 | 491.967 | 134.972 | 118.441 | 77.020 | 63.837 | 51.293 |
| Memory usage (MiB, GPU) | 1107.820 | 538.897 | 402.478 | 323.974 | 286.282 | 212.643 | 183.501 |

(a) Sample C (slice 2 and slice 3) of the DLPFC dataset.

| $dx$ | 10 | 20 | 30 | 40 | 50 | 70 | 100 |
|---|---|---|---|---|---|---|---|
| Mean scores | 0.469 | 0.472 | 0.471 | 0.470 | 0.468 | 0.456 | 0.446 |
| Runtime (s) | 190.985 | 72.151 | 54.094 | 51.031 | 50.468 | 52.081 | 50.597 |
| Memory usage (MiB, GPU) | 168.626 | 59.361 | 40.101 | 38.139 | 34.557 | 32.890 | 32.733 |

(b) Mouse liver map. Mean scores indicate the average cosine similarity scores of selected 136 liver marker genes.

the estimated transformation against moderate changes in discretisation granularity. As a result, the recovered diffeomorphic mapping remains largely consistent across practical resolution settings, even though the raster grid varies. In contrast, computational cost increases substantially as $dx$ decreases, as reflected in both runtime and GPU memory usage.

Considering both computational efficiency and alignment performance, we adopt $dx \approx 1$ unit for spot-level data and $dx = 30\,\mu$m for cell-level data as default settings, which provide a favourable balance between spatial fidelity and resource usage without requiring fine-grained hyper-parameter tuning.

###### S2.1.4 Robustness to spatial overlap and incomplete coverage of informative genes

Methodologically, informative genes are ranked according to their predictability from local spatial context, and this selection process does not explicitly enforce complementary spatial coverage. To evaluate the robustness of GALA to situations where informative genes exhibit substantial spatial overlap or incomplete tissue coverage, we selected alternative genes from the top 10 informative genes after rasterization in the MBSP dataset (Fig. S6) and assessed the alignment performance under these conditions. The results were compared with the default informative genes used in the manuscript.

Specifically, we constructed two sets of partially distributed expressed genes with $p = 3$. The first set consists of *Nrgn*, *Ddn*, and *Camk2a* (highlighted by orange rectangles in Fig. S6). These genes are predominantly expressed in excitatory neurons within cortical and hippocampal layers and are mainly concentrated on the left side of each tensor image. The second set includes *Pcp2*, *Cbln3*, and *Pvalb* (highlighted by green rectangles in Fig. S6). These genes are spatially distributed across cortical layers and are mainly concentrated on the right side of each tensor image. Together, these gene sets exhibit noticeable spatial overlap and do not collectively cover the entire tissue section. When replacing these genes with the top three informative genes used in the manuscript, the spatial cross-correlation (SCC) scores remain nearly unchanged (left-biased: 0.666; right-based: 0.662; original setting: 0.669 as shown in Fig. S7). This result demonstrates that GALA's alignment performance is not highly sensitive to the exact choice of informative genes, even when their spatial patterns overlap or fail to fully cover the tissue.

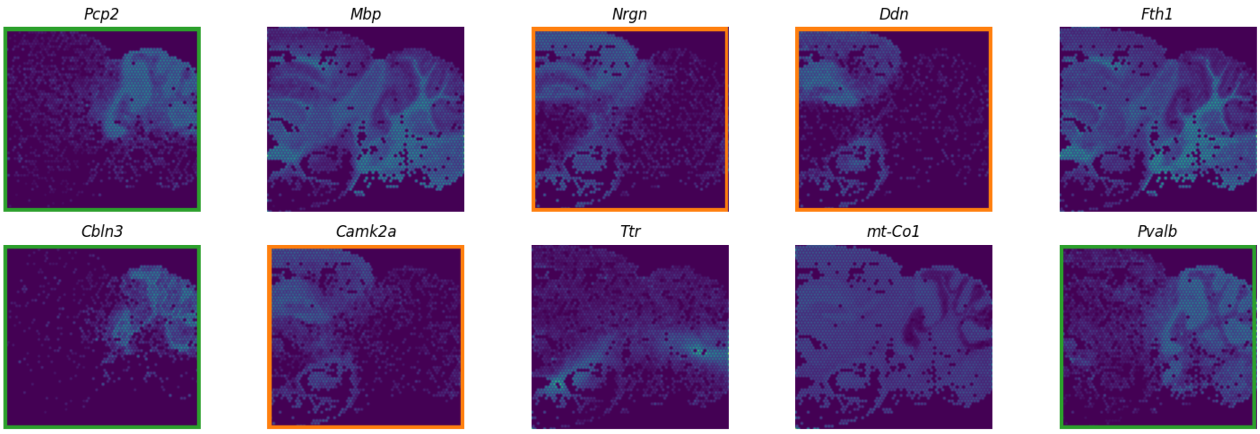

Figure S6: **The top ten informative genes after rasterization.**

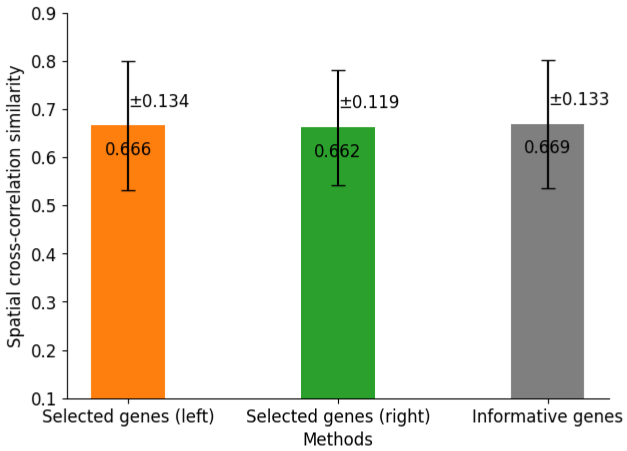

Figure S7: **SCC scores under different informative gene selections (mean ± s.d.)**. "Selected genes (left)" refers to *Nrgn*, *Ddn*, and *Camk2a*. "Selected genes (right)" refers to *Pcp2*, *Cbln3*, and *Pvalb*. "Informative genes" refers to the top three informative genes used in the manuscript.

#### S2.2 Transformation properties of GALA

As described in foundational works on the LDDMM framework(Beg et al., 2005) and its applications (Tward et al., 2020; Clifton et al., 2023), LDDMM provides a mathematically rigorous formulation for modelling non-linear spatial transformations. Its key properties directly motivate the design choices in GALA.

##### S2.2.1 Invertibility and bidirectional consistency

By construction, the deformation $\phi^v$ generated by LDDMM is diffeomorphic and therefore invertible, smooth, and topology-preserving. Since the affine transformation $A$ used in GALA is also invertible, their composition $(\phi^v \cdot A)$ remains invertible and continuous. As a result, forward (source-to-target) and backward (target-to-source) alignments are theoretically well-defined.

To empirically verify this property, we performed additional experiments by swapping source and target roles in the primary within-resolution alignment tasks, including DLPFC Sample C (Slice 2 $\leftrightarrow$ Slice 3, spot-level), partial-overlap experiments (Slice 2 $\leftrightarrow$ partial Slice 3, spot-level), and mouse liver replicates (cell-level). In practice, numerical discretisation, optimisation asymmetry, and intensity modelling may introduce small discrepancies, so the two directions are not exactly identical but are expected to be highly consistent. As shown in Table S3, alignment accuracy remains highly consistent between forward and reverse directions across all datasets.

Table S3: Forward and reverse alignment consistency analysis

| Label consistency | Accuracy |
|---|---|
| Slice 3 $\rightarrow$ Slice 2 | 0.856 |
| Slice 2 $\rightarrow$ Slice 3 | 0.847 |

(a) Forward and reverse alignment on DLPFC Sample C (spot-level).

| Label consistency | Accuracy |
|---|---|
| Partial Slice 3 $\rightarrow$ Slice 2 | 0.874 |
| Slice 2 $\rightarrow$ Partial Slice 3 | 0.863 |

(b) Forward and reverse alignment under partial overlap.

| Cosine similarity | Average scores |
|---|---|
| Replicate 2 $\rightarrow$ Replicate 1 | 0.471 |
| Replicate 1 $\rightarrow$ Replicate 2 | 0.477 |

(c) Forward and reverse alignment on mouse liver replicates (cell-level). Cosine similarity scores indicates the median scores of 136 liver marker genes.

For a more direct geometric verification, we evaluated the deviation from inversion on DLPFC Sample C. Let $(\phi^v \cdot A)_{3\to2}$ denote the transformation from Slice 3 to Slice 2, and $(\phi^v \cdot A)_{2\to3}$ denote the reverse mapping. Here, '$\cdot$' denotes function composition, i.e., $(\phi^v \cdot A)(\boldsymbol{x}) = \phi^v(A(\boldsymbol{x}))$. Ideally, the forward transformation should be the inverse of the reverse transformation, meaning

$$(\phi^v \cdot A)_{2\to3}^{-1} \cdot (\phi^v \cdot A)_{3\to2} = \mathrm{Id},$$

where Id denotes the identity mapping. We computed the pointwise deviation

$$\|(\phi^v \cdot A)_{2\to3}^{-1} \cdot (\phi^v \cdot A)_{3\to2} - \mathrm{Id}\|,$$

and report the mean error magnitude as 6.331 (in the native spatial coordinate unit). Fig. S8 visualises both forward and the inverse of reverse transformations and the spatial distribution of inversion error. Given that tissue sections span several thousand spatial units, this deviation represents a small fraction of the global spatial extent, confirming strong directional consistency in practice.

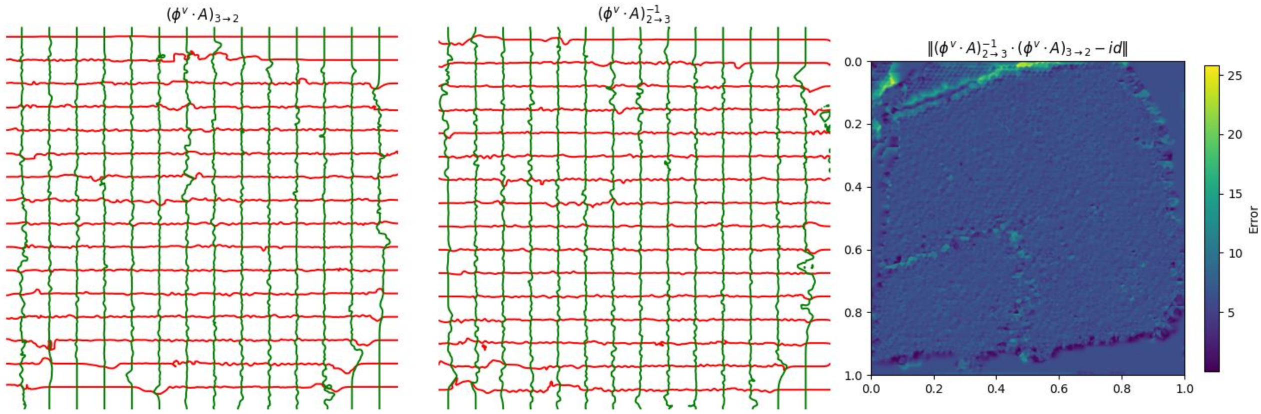

Figure S8: Forward and reverse transformations of DLPFC Sample C (Slice 2 $\leftrightarrow$ 3). The left panel shows the forward transformation $(\phi^v \cdot A)_{3\to2}$, the middle panel shows the reverse transformation $(\phi^v \cdot A)_{2\to3}$, and the right panel displays the inversion error $\|(\phi^v \cdot A)_{2\to3}^{-1} \cdot (\phi^v \cdot A)_{3\to2} - \mathrm{Id}\|$, with a mean error of 6.331.

We note that this bidirectional consistency analysis applies primarily to within-resolution alignments. For cross-resolution or cross-modality alignment (e.g., transcriptomics–histology), the optimisation problem becomes intrinsically asymmetric due to differences in spatial sampling density and modality-specific signal characteristics. In such cases, aligning the higher-resolution modality toward the lower-resolution reference is generally more numerically stable, and exact inversion symmetry is not expected. In fact, the reverse direction involves reconstructing fine-scale structure from coarse measurements, constituting an ill-posed inverse problem closely related to cell deconvolution.

##### S2.2.2 Large deformation and synthetic validation

LDDMM parameterises spatial transformations through time-dependent velocity fields $v_t$, generating diffeomorphic flows via integration over time as described in Eqn. (S1). Unlike small-displacement linear elastic or spline-based models, this formulation does not assume infinitesimal deformation. Instead, spatial displacement is accumulated through integration of a smooth velocity field, allowing substantial non-linear geometric warping while preserving topology. This is the origin of the term "Large Deformation" in LDDMM.

However, LDDMM assumes that the underlying transformation between sections can be represented as a smooth diffeomorphism. Deformations that introduce discontinuities or topological changes violate this modelling assumption and fall outside its theoretical regime.

To empirically validate the large-deformation capability, we constructed synthetic deformation fields on the DLPFC dataset (Sample C, Slice 2) and aligned the resulting deformed slices back to the original slice using GALA. Let $\xi_x(x)$ and $\xi_y(y)$ denote independent Gaussian white noise fields sampled on a regular grid. We define the displacement field

$$u(\boldsymbol{x}) = (u_x(x), u_y(y)),$$

with

$$u_x(x) = \lambda(G_\sigma * \xi_x)(x), \quad u_y(y) = \lambda(G_\sigma * \xi_y)(y),$$

where $*$ denotes convolution and

$$G_\sigma(x) = \frac{1}{2\pi\sigma^2} \exp\left(-\frac{\|x\|^2}{2\sigma^2}\right)$$

is a Gaussian kernel with bandwidth $\sigma$. The bold $\boldsymbol{x} = (x, y)$ denotes the gird location in Cartesian coordinates. The scalar $\lambda$ controls the deformation magnitude, while $\sigma$ determines the spatial smoothness of the displacement field. The deformed slice is obtained as

$$\boldsymbol{x}' = \boldsymbol{x} + u(\boldsymbol{x}),$$

and subsequently aligned back to the original target slice using GALA to recover spatial correspondence.

We first fixed $\lambda = 100$ and varied $\sigma \in \{1, 2, 3, 4, 5\}$ to assess the impact of smoothness as shown in Fig. S9a. Before alignment, the label consistency accuracy between the deformed slice and the original slice increased with $\sigma$: 0.647, 0.787, 0.822, 0.850, and 0.867, respectively. After applying GALA, accuracies improved to 0.642, 0.799, 0.835, 0.859, and 0.878. This indicates that smoother, large-scale warping (larger $\sigma$) is more accurately recovered, whereas highly irregular local distortions (small $\sigma$) are intrinsically harder to align due to high-frequency components.

Next, we fixed $\sigma = 3$ and varied $\lambda \in \{50, 75, 100, 125, 150\}$ to evaluate the effect of deformation magnitude as shown in Fig. S9b. Before alignment, label consistency decreased from 0.894 to 0.754 as $\lambda$ increased, reflecting increasing difficulty of aligning more severely deformed slices. After applying GALA, the corresponding accuracies increased to 0.901, 0.864, 0.829, 0.815, and 0.789, showing that GALA effectively recovers spatial correspondence even under large-amplitude smooth deformations.

These results demonstrate that GALA can robustly recover spatial correspondence under substantial smooth non-linear deformations. Alignment accuracy is strongly influenced by the smoothness of the deformation: highly irregular local distortions (small $\sigma$) are intrinsically more difficult to recover, reflecting the high smoothness requirement imposed by the LDDMM framework. In contrast, large-amplitude but smooth deformations (large $\lambda$ with moderate to large $\sigma$) are handled effectively, indicating that GALA's main limitation lies in dealing with high-frequency, irregular distortions rather than overall deformation magnitude.

To further evaluate the robustness of the GA-based affine initialization under non-rigid transformations, we performed an additional experiment in which the deformed slices were first rotated by $\theta = 30$ before applying the synthetic smooth non-linear displacements (Fig. S9c). Using $\lambda = 100$ and $\sigma \in \{1, 2, 3, 4, 5\}$, the pre-alignment label consistency accuracies were 0.450, 0.491, 0.496, 0.500, and 0.509, reflecting the combined effect of global rotation and non-linear deformation. After applying GALA, the accuracies improved to 0.633, 0.779, 0.825,

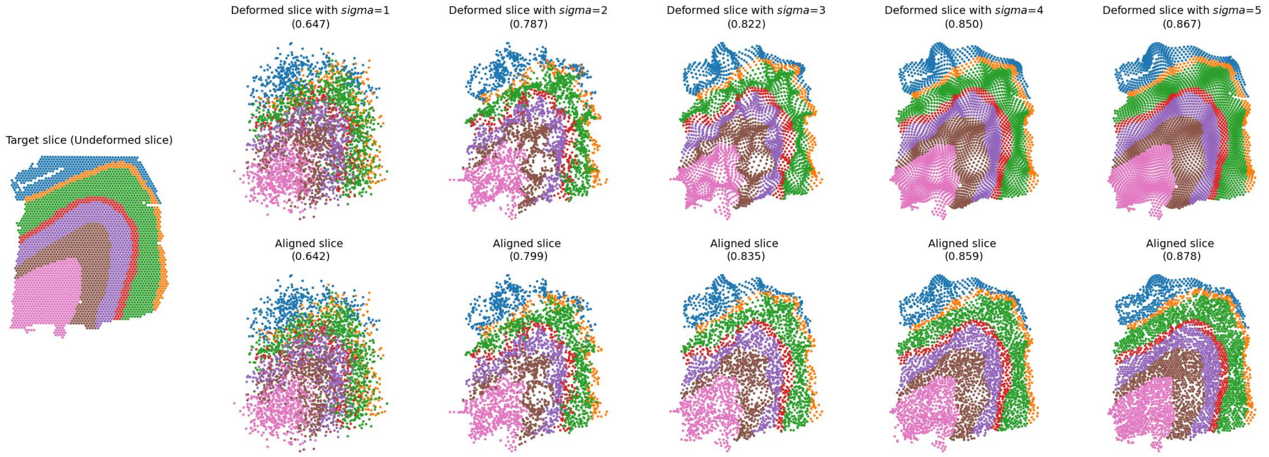

(a) Deformed slices across $\sigma = \{1, 2, 3, 4, 5\}$ with fixed $\lambda = 100$ and aligned slices after GALA.

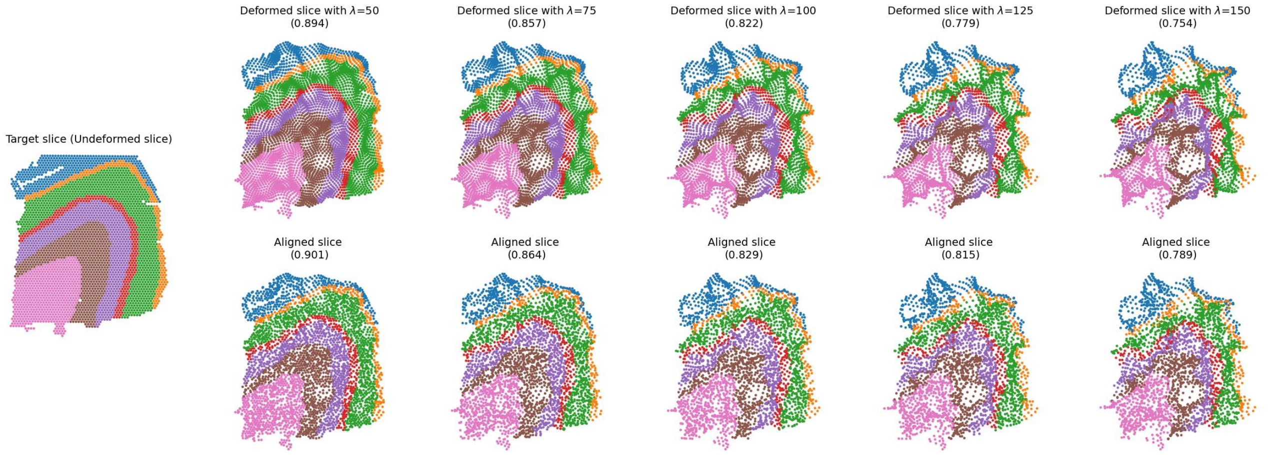

(b) Deformed slices across $\lambda = \{50, 75, 100, 125, 150\}$ with fixed $\sigma = 3$ and aligned slices after GALA.

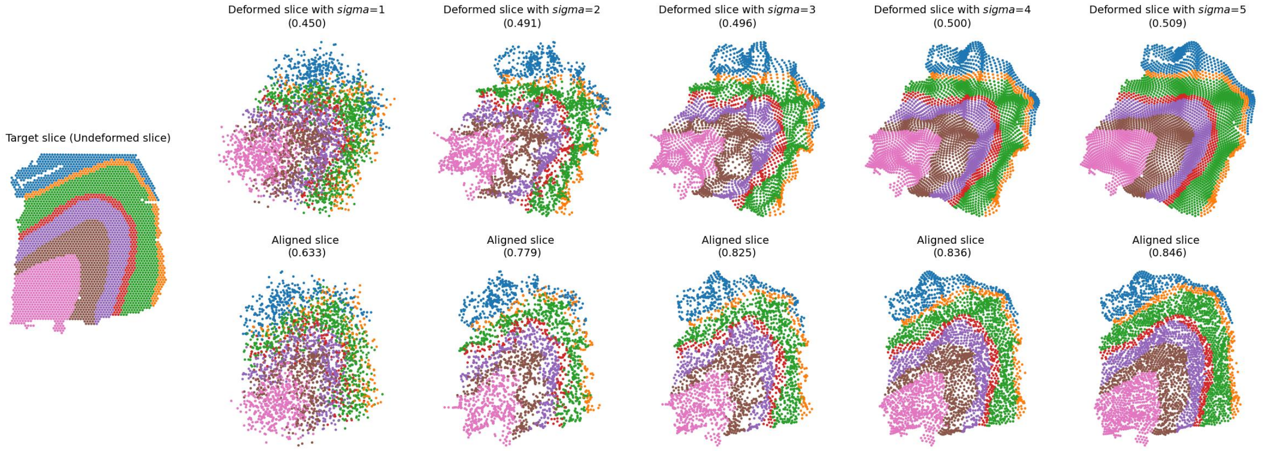

(c) Rotated and deformed slices across $\theta = 30, \sigma = \{1, 2, 3, 4, 5\}$ with fixed $\lambda = 100$ and aligned slices after GALA.

Figure S9: Synthetic validation of GALA under large non-linear deformations. The experiments systematically evaluate the effects of spatial smoothness ($\sigma$), deformation magnitude ($\lambda$), and additional global rotation ($\theta = 30$) on alignment performance. Values in brackets denote label consistency accuracy. Spots are annotated with six neocortical layers (Layer 1–6) and white matter (WM).

0.836, and 0.846. Notably, even for small $\sigma$ values ($\sigma = 1, 2$), substantial improvement is observed, indicating that the affine transformation is correctly estimated despite the presence of rotation and irregular deformation. As $\sigma$ increases ($\sigma \geq 3$), alignment accuracy further improves and approaches that of the non-rotated experiments, suggesting that performance is primarily governed by deformation smoothness rather than failure of the affine initialization.

To further evaluate the robustness of the GA-based affine initialization under non-rigid transformations, we performed an additional experiment in which the deformed slices were first rotated by $\theta = 30$ before applying the synthetic smooth non-linear displacements (Fig. S9c). Using a large deformation amplitude ($\lambda = 100$), the pre-alignment label consistency was low (0.450–0.509), reflecting substantial global misalignment. After applying GALA, post-alignment accuracies improved substantially (0.633–0.846), indicating that the GA-based affine initialization successfully recovers the global transformation despite the rotation and non-rigid deformation. The post-alignment accuracy is slightly lower than in the non-rotated experiments (see Fig. S9a) because the global rotation perturbs local neighborhoods, limiting the precision of LDDMM refinement. Overall, these results demonstrate that the GA-based affine stage reliably estimates global transformations and provides a solid initialization for local LDDMM refinement, while minor reductions in accuracy are primarily due to challenges in refining non-smooth local deformations rather than failure of the affine initialization.

Taken together, these synthetic experiments delineate the deformation regime in which GALA operates reliably. Across variations in smoothness ($\sigma$), deformation magnitude ($\lambda$), and additional global rotation, alignment performance is primarily governed by the smoothness of the underlying transformation. Highly irregular, high-frequency distortions (small $\sigma$) remain intrinsically challenging, whereas smoother deformations are consistently recovered with high accuracy, even at large amplitudes. Importantly, the additional rotation experiments demonstrate that the GA-based affine initialization remains stable under combined global and non-linear transformations, and does not constitute the primary limiting factor. Overall, these results indicate that GALA is robust to substantial smooth non-linear deformations and moderate global misalignment, with performance chiefly determined by deformation regularity rather than displacement magnitude or affine estimation stability.

#### S2.3 Computational performance benchmarking

All experiments were conducted on a Linux workstation (kernel 6.11.0-26-generic) equipped with an Intel(R) Core(TM) i7-14700K CPU (28 cores), 62.6 GB RAM, and an NVIDIA GeForce RTX4090D GPU. GALA was implemented in PyTorch 2.6.0 with CUDA 12.4.

We benchmarked GALA against existing methods across different datasets. Each experiment was repeated three times, and the average runtime as well as CPU and GPU memory consumption are reported in Tab. S4. Methods supporting GPU acceleration were executed on the same GPU as GALA, while CPU-only methods were run on the same CPU, ensuring fair hardware conditions.

For DLPFC slices 2 and 3 using transcriptomics only (Tab. S4a), GALA completed alignment in 135.22 s, faster than the non-rigid method GPSA (245.61 s) and the coarse-to-fine method ST-GEARS (154.69 s). Its memory usage (420.43 MiB CPU, 279.80 MiB GPU) remained moderate compared with several baselines, some of which exhibited substantially higher memory consumption.

In partial alignment (Tab. S4b), GALA required 97.03 s with 417.03 MiB CPU and 167.00

MiB GPU memory, substantially outperforming PASTE2, which required significantly longer runtime and higher CPU memory usage.

On MBSP sections (Tab. S4c), GALA was faster than GPSA (155.41 s vs 248.28 s) while maintaining comparable GPU memory usage and lower CPU memory consumption.

For the MERFISH mouse liver dataset (Tab. S4d), several methods encountered scalability limitations due to excessive memory demands ( 500 GB). Although SLAT completed its training phase quickly, it required nearly two hours for the subsequent graph-based cell matching. Both STalign and GALA employ LDDMM for local diffeomorphic deformation; however, GALA completed the alignment in 54.11 s compared with 123.17 s for STalign. In addition to the shorter runtime, GALA required lower memory usage (394.86 MiB CPU and 30.03 MiB GPU) than STalign (528.96 MiB CPU and 71.48 MiB GPU), reflecting its more streamlined optimisation and reduced parameters.

In the mismatched-resolution scenario (Tab. S4e), GALA aligned 149,023 cells to 2,634 spots in 73.30 s, comparable to SLAT (71.78 s), while using substantially less GPU memory (16.73 MiB vs 3363.43 MiB).

For transcriptomics–histology alignment (Tab. S4f, g), GALA consistently outperformed STalign in runtime while maintaining lower GPU memory usage across both lung and breast cancer datasets.

Table S4: Comparison of computation time between GALA and other baselines

| Methods | PASTE | STAligner | GPSA | SLAT | ST-GEARS | GALA |
|---|---|---|---|---|---|---|
| Average Runtime (s) | 21.02 | 8.75 | 245.61 | 4.05 | 154.69 | 135.22 |
| Average CPU Memory (MiB) | 512.66 | 719.14 | 514.09 | 1206.08 | 244.13 | 420.43 |
| Average GPU Memory (MiB) | 700.33 | 448.14 | 850.26 | 145.43 | 0 | 279.80 |

(a) Slice 2 and 3 (3,566 and 3,635 spots, 10,629 shared genes) from Sample C in DLPFC dataset.

| Methods | PASTE2 | GALA |
|---|---|---|
| Average Runtime (s) | 359.53 + 960.26 | 97.03 |
| Average CPU Memory (MiB) | 996.04 | 417.03 |
| Average GPU Memory (MiB) | 0 | 167.00 |

(b) Partial alignment (60% leftmost retained) between slices 2 and 3 from Sample C in DLPFC dataset.

| Methods | GPSA | GALA |
|---|---|---|
| Average Runtime (s) | 248.28 | 155.41 |
| Average CPU Memory (MiB) | 476.98 | 183.71 |
| Average GPU Memory (MiB) | 233.72 | 248.15 |

(c) MBSP sections using molecular and morphology information (2,805 and 284 spots, 4,260 shared genes).

| Methods | STalign | SLAT | GALA |
|---|---|---|---|
| Average Runtime (s) | 123.17 | 3600+ | 54.11 |
| Average CPU Memory (MiB) | 528.96 | 2891.89 | 394.86 |
| Average GPU Memory (MiB) | 71.48 | 8952.55 | 30.03 |

(d) Mouse liver sections (300,038 and 310,932 cells, 347 shared genes).

| Methods | SLAT | GALA |
|---|---|---|
| Average Runtime (s) | 71.78 | 73.30 |
| Average CPU Memory (MiB) | 708.18 | 371.95 |
| Average GPU Memory (MiB) | 3363.43 | 16.73 |

(e) Mouse brain sections (149,023 cells to 2,634 spots, 248 shared genes).

| Methods | STalign | GALA |
|---|---|---|
| Average Runtime (s) | 317.28 | 90.32 |
| Average CPU Memory (MiB) | 498.56 | 406.87 |
| Average GPU Memory (MiB) | 291.21 | 89.50 |

(f) FFPR lung cancer sections (image: $2051 \times 2759$ pixels, transcriptomic coordinates: $5475 \times 7521$).

| Methods | STalign | GALA |
|---|---|---|
| Average Runtime (s) | 512.06 | 74.16 |
| Average CPU Memory (MiB) | 438.55 | 430.06 |
| Average GPU Memory (MiB) | 132.20 | 48.18 |

(g) FFPR breast cancer sections (image: $4345 \times 2508$ pixels, transcriptomic range: $11441 \times 7093$).

### S3 Supplementary results

#### S3.1 Supplementary tables

Table S5: Evaluation of label consistency accuracy, ARI, and NMI for Sample C across all alignment baselines. ‘Raw’ denotes the unaligned slices.

| Metric | Slice | Raw | PASTE | STAligner | GPSA | SLAT | ST-GEARS | GALA |
|---|---|---|---|---|---|---|---|---|
| Accuracy | $2 \rightarrow 1$ | 0.786 | **0.876** | 0.753 | 0.875 | 0.871 | 0.869 | 0.872 |
| | $3 \rightarrow 2$ | 0.688 | 0.827 | 0.599 | 0.860 | 0.827 | 0.816 | **0.867** |
| | $4 \rightarrow 3$ | 0.776 | 0.815 | 0.841 | 0.764 | 0.782 | 0.791 | **0.864** |
| ARI | $2 \rightarrow 1$ | 0.369 | 0.370 | 0.369 | 0.368 | 0.371 | 0.387 | **0.395** |
| | $3 \rightarrow 2$ | 0.303 | 0.322 | **0.395** | 0.379 | 0.383 | 0.394 | 0.392 |
| | $4 \rightarrow 3$ | 0.331 | 0.268 | 0.386 | 0.343 | 0.360 | 0.289 | **0.399** |
| NMI | $2 \rightarrow 1$ | 0.495 | 0.497 | 0.506 | 0.494 | 0.497 | 0.503 | **0.508** |
| | $3 \rightarrow 2$ | 0.431 | 0.445 | 0.486 | **0.503** | 0.505 | 0.497 | 0.494 |
| | $4 \rightarrow 3$ | 0.453 | 0.441 | 0.480 | 0.456 | 0.478 | 0.441 | **0.521** |

#### S3.2 Supplementary figures

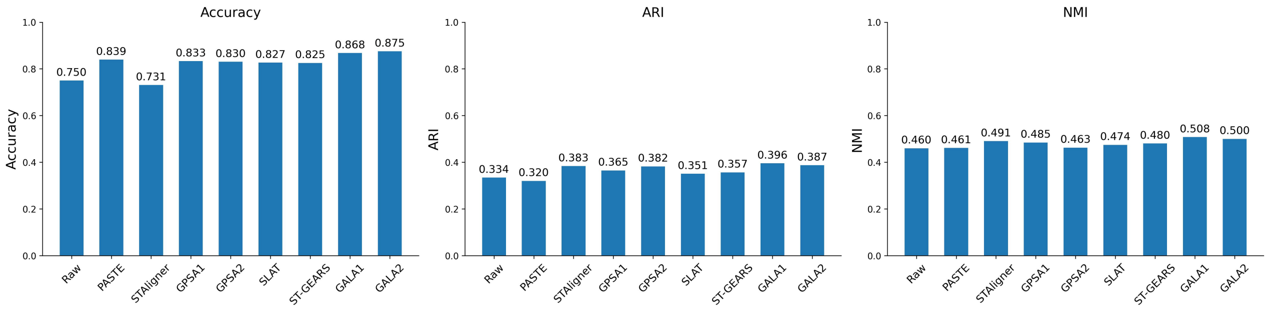

(a) Mean quantitative scores of aligned slices in Sample C.

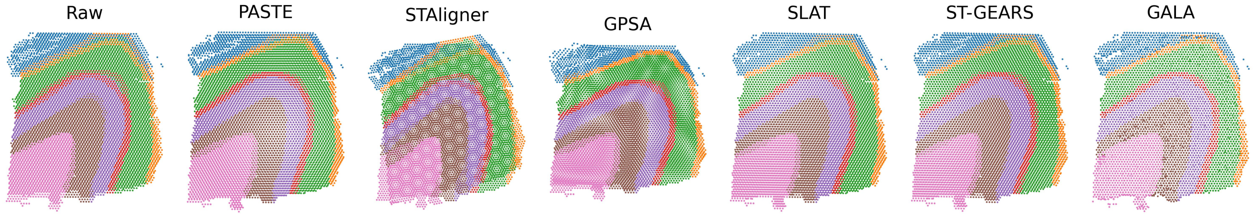

(b) Stacked slices of slice 1 and aligned slice 2

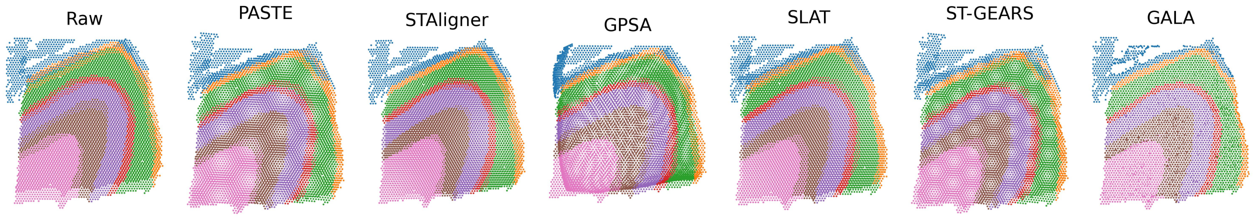

(c) Stacked slices of slice 3 and aligned slice 4

Figure S10: **Supplementary alignment results for Fig. 2**.

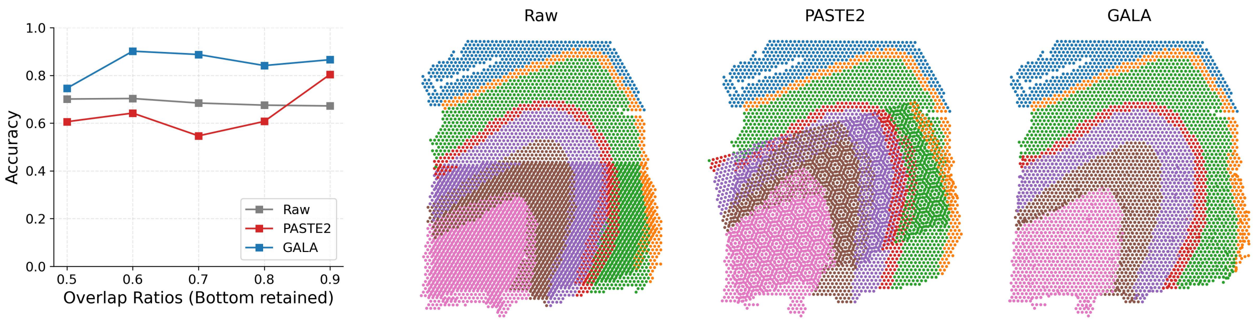

(a) Supplementary results for Fig. 3b.

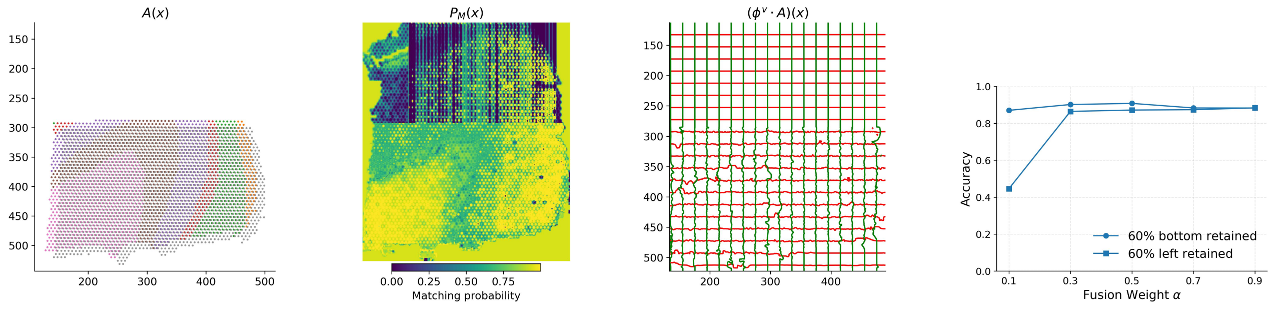

(b) Supplementary results for Fig. 3c.

(c) $\alpha$-Sensitivity analysis

Figure S11: **Supplementary results for partial alignment**. (a) Label consistency accuracy when aligning the cropped source data (bottom retained slice 3) to the target data (slice 2) across different overlap ratios and baseline methods, alongside visual comparisons for the partial alignment at an overlap ratio of 0.6. (b) Illustration of GALA's alignment mechanism for the 'bottom 60% retained' scenario, showing the learned estimators $A(\boldsymbol{x})$ for the global affine transformation, $P_M(\boldsymbol{x})$ for matching probabilities within the overlapping region, and the local diffeomorphic mapping $(\phi^v \cdot A)(\boldsymbol{x})$. (c) Sensitivity analysis of the parameter $\alpha$, evaluated across two types of partial alignment and a range of overlap ratios.

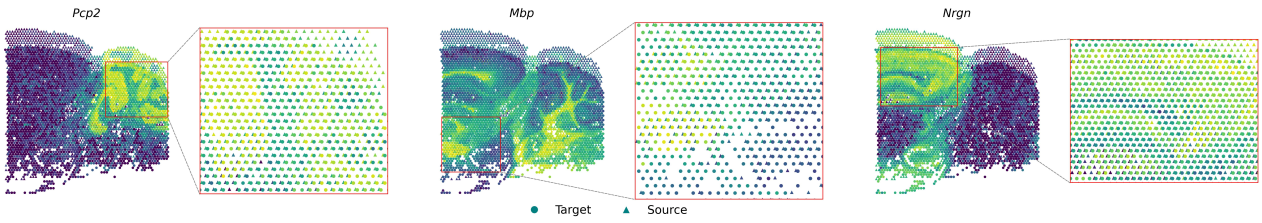

Figure S12: **Supplementary results for Fig. 4d.** Raw stacked slices of three informative genes, coloured according to gene expression levels.

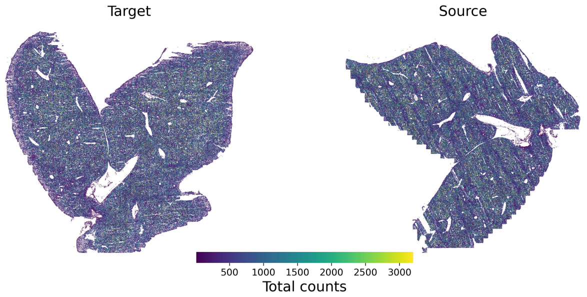

(a) Supplementary results for Fig. 5a.

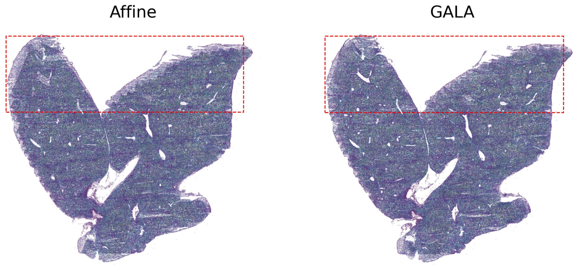

(b) Supplementary results for Fig. 5c.

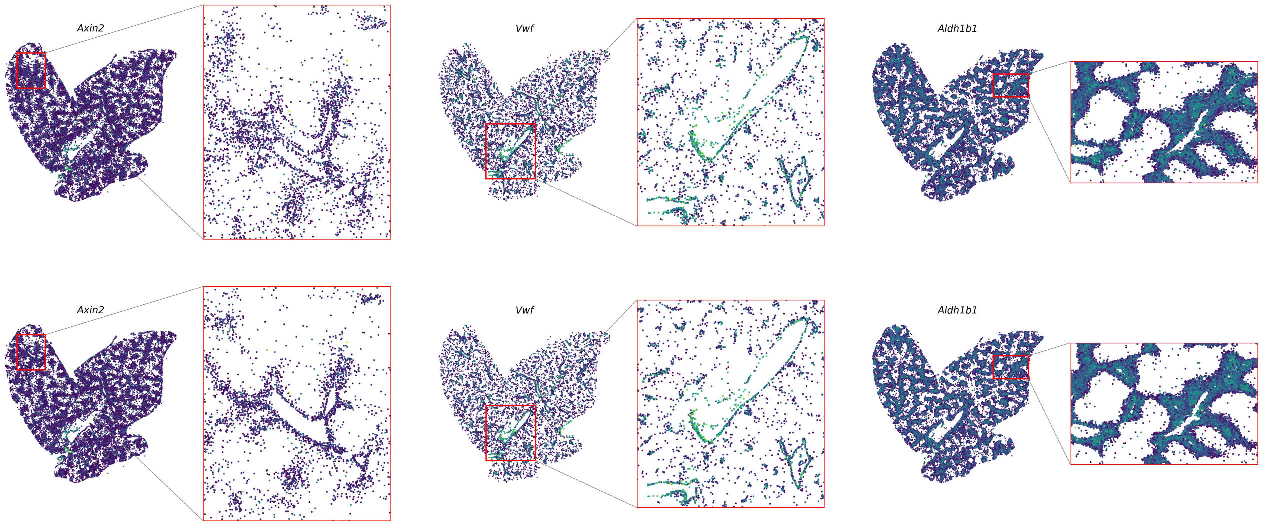

(c) Supplementary results for Fig. 5d.

Figure S13: **Supplementary results for Fig.5 coloured by gene expression.** Brighter colours indicate higher total gene expression.

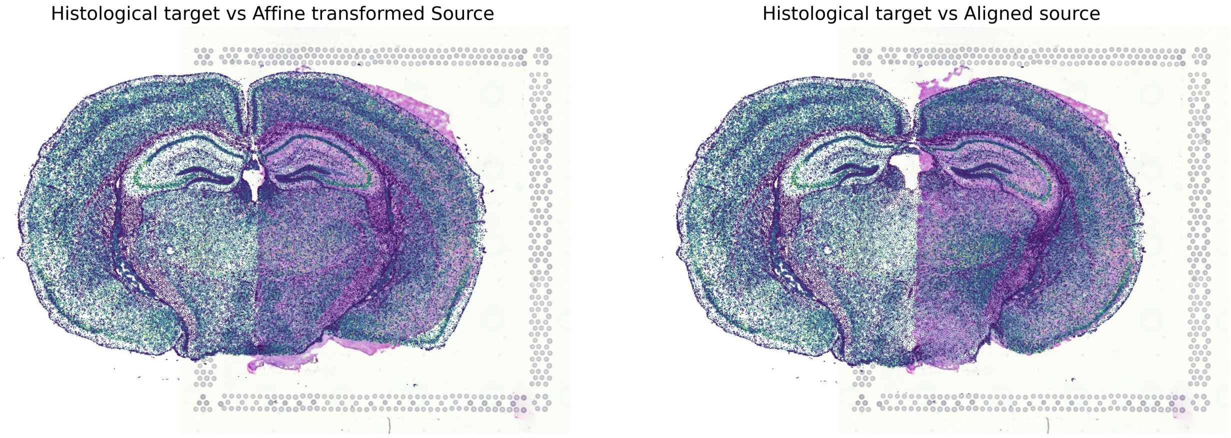

Figure S14: Overlaid aligned slices of cell-resolved data on the target histological slice in the left hemisphere: Affine-aligned (left) versus GALA-refined alignment (right).

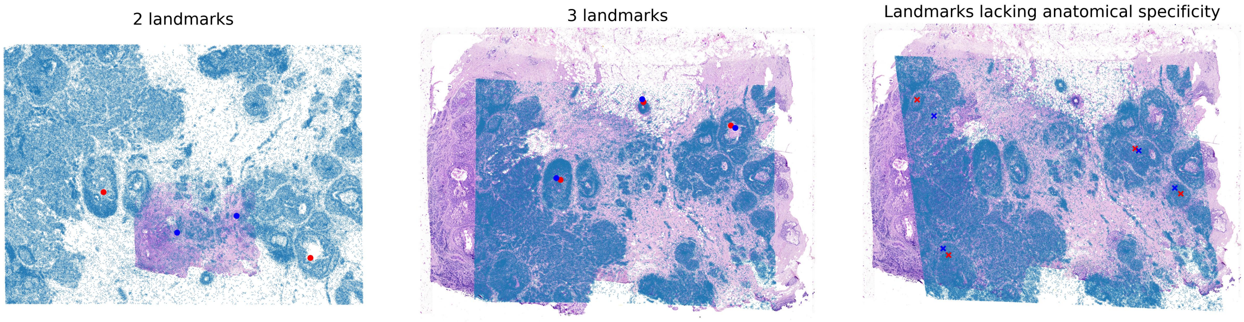

Figure S15: **Evaluation of pre-defined landmarks in STalign.** The first two panels show alignment results using different numbers of landmarks. The last panel shows alignment based on randomly chosen landmarks lacking anatomical specificity.

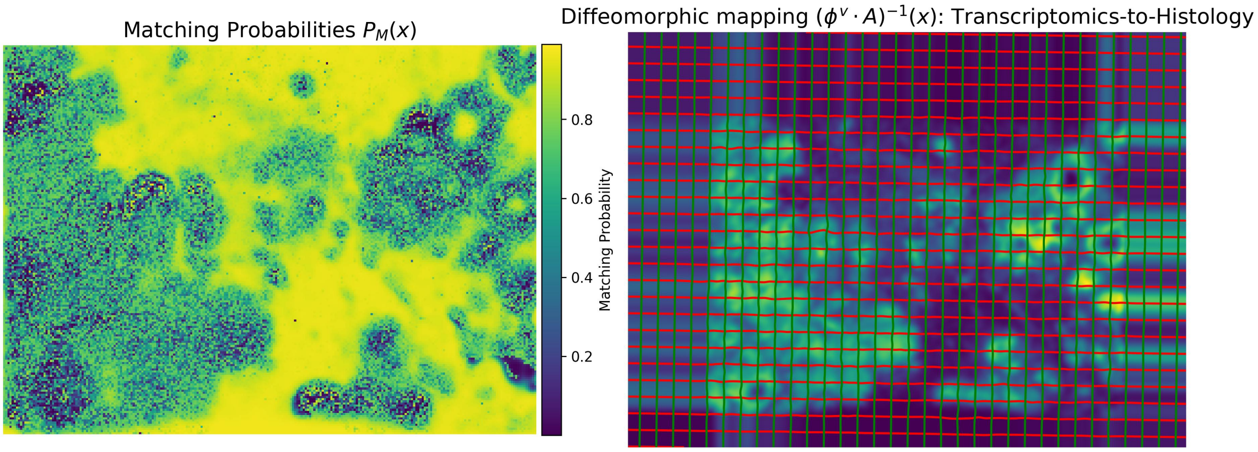

Figure S16: $P_M(\boldsymbol{x})$ **and** $(\phi^v \cdot A)^{-1}(\boldsymbol{x})$. The background of right panel shows the rasterised total expression of the target. In this example, the processed histological image is aligned as the source to the target using molecular information. Accordingly, the final result requires computing the inverse of $(\phi^v \cdot A)(\boldsymbol{x})$.
